# Inferring cell-cell interactions from pseudotime ordering of scRNA-Seq data

**DOI:** 10.1101/2021.07.28.454054

**Authors:** Dongshunyi Li, Jeremy J. Velazquez, Jun Ding, Joshua Hislop, Mo R. Ebrahimkhani, Ziv Bar-Joseph

## Abstract

A major advantage of single cell RNA-Sequencing (scRNA-Seq) data is the ability to reconstruct continuous ordering and trajectories for cells. To date, such ordering was mainly used to group cells and to infer interactions within cells. Here we present TraSig, a computational method for improving the inference of cell-cell interactions in scRNA-Seq studies. Unlike prior methods that only focus on the average expression levels of genes in clusters or cell types, TraSig fully utilizes the dynamic information to identify significant ligand-receptor pairs with similar trajectories, which in turn are used to score interacting cell clusters. We applied TraSig to several scRNA-Seq datasets. As we show, using the ordering information allows TraSig to obtain unique predictions that improve upon those identified by prior methods. Functional experiments validate the ability of TraSig to identify novel signaling interactions that impact vascular development in liver organoid.

## Background

The ability to profile cells at the single cell level enabled the identification of new cell types and additional markers for known cell types as well as the reconstruction of cell type specific regulatory networks [1, 2]. Several methods have been developed to group or cluster cells in scRNA-Seq data [3] and to reconstruct trajectories and pseudotime for time series scRNA-Seq data [4]. Such methods have mainly focused on the expression similarity between cells in the same cluster or at consecutive time points and on the differences in transcriptional regulation between cell types and over time [5]. More recently, a number of methods have been developed to infer another type of interaction from scRNA-Seq data: signaling between cell clusters or cell types [6]. These methods attempt to identify ligands in one of the clusters or cell types and corresponding receptors in another cluster and then infer interactions based on the average expression of these ligand-receptor pairs. For example, CellPhoneDB [7] scores ligand-receptor pairs using their mean expression values in two clusters and assigns significance levels using permutations tests. SingleCellSingleR[8] designs a score based on the product of ligand-receptors’ mean expression values in two clusters and selects ligand-receptors scoring above a predefined threshold.

While successful, most current methods for inferring cell-cell interactions from scRNA-Seq data only use the average expression levels of ligands and receptors in the two clusters or cell types they test [6]. While this may be fine for steady state populations (for example, different cell types in adult tissues), for studies that focus on development or response modeling, such averages do not take full advantage of the available data in scRNA-Seq studies. Indeed, even cells on the same branch are often ordered in such studies using various pseudotime ordering methods [9]. In such cases, cells on the same branch (or cluster) cannot be assumed to be homogeneous with respect to the expression of key genes. Using average analysis for such clusters may lead to inaccurate predictions about the relationship between ligands and receptors in two different (though parallel in terms of timing) branches. Specifically, Figure 1 presents four cases of pseudotime orderings for a ligand and its corresponding receptor in two different branches. While the *average* expression of a ligand and receptor in two different branches are the same, the first two cases are unlikely to strongly support an interaction between these two cell types while the third and fourth, where both are either increasing or decreasing in their respective ordering, are much more likely to hint at real interactions between the groups. In other words, if two groups of cells are interacting, then we expect to see the genes encoding signaling molecules in these groups co-express at a similar pace along the pseudotime.

**Figure 1:**
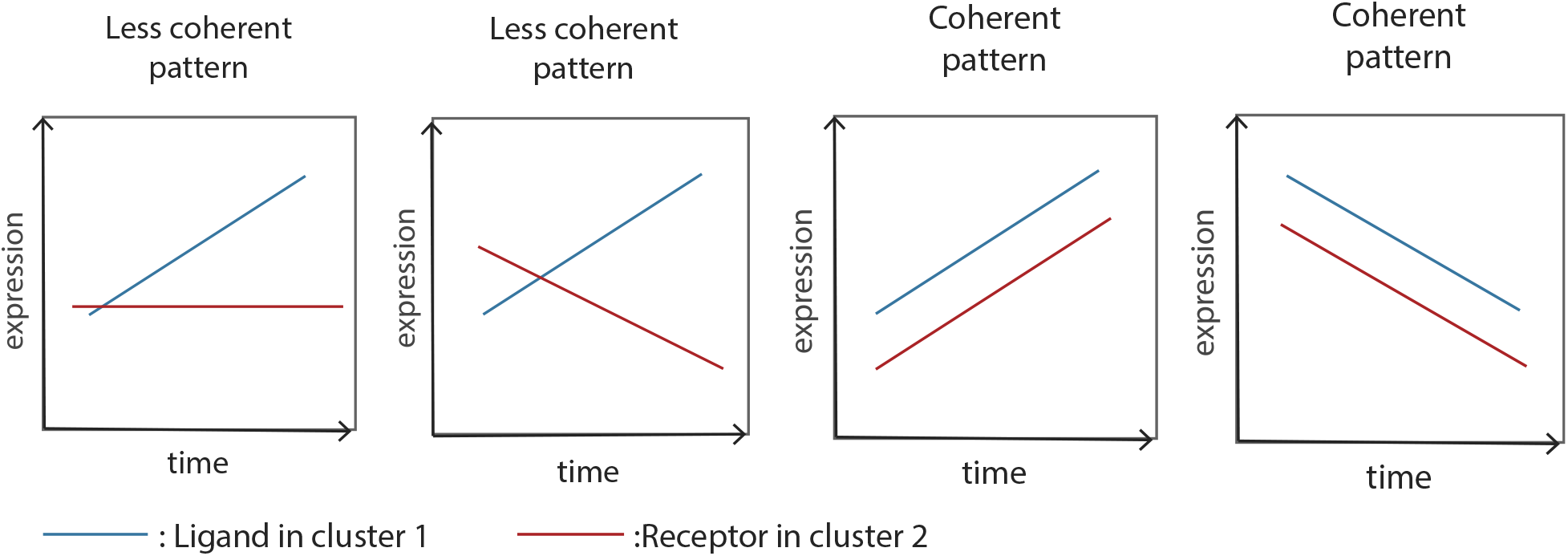
Example cases where the *average* expressions of the ligand and receptor that are known to interact are the same. Of these four figures only the last two represent correlated activation and repression of these proteins. Methods that only use the average expression of genes in clusters cannot differentiate between these 4 profiles and so will score all of them the same.

To enable the use of pseudotime ordering for predicting cell type interactions between dynamically changing cell populations, we developed TraSig. TraSig can use several of the most popular pseudotime ordering and trajectory inference methods to extract expression patterns for ligands and receptors in different edges of the trajectory using a sliding window approach. It then uses these profiles to score temporal interactions between ligands and their known receptors in different edges corresponding to the same time. Permutation testing is used to assign significance levels to specific pairwise interactions and scores are combined to identify significant cluster-cluster interactions.

We applied TraSig to a number of scRNA-Seq datasets and compared its performance to a number of popular methods for inferring signaling interactions from scRNA-Seq data. As we show, the ability to utilize the temporal information in the analysis improves the accuracy of predicted relevant pairs and leads to distinct predictions that are not identified by other methods that rely on average expression. We experimentally validated a number of interaction predictions from TraSig for liver organoid differentiation data.

## Results

We developed a computational method, TraSig for inferring cell-cell interactions from pseudotime ordered data. Figure 2 presents an overview of the method. We start by using a trajectory inference method to obtain grouping and pseudotime ordering for cells in the dataset. Here we use Continuous-state Hidden Markov Model (CSHMM) [10] for this, though as discussed below, TraSig can be applied to results from other pseudotime ordering methods. We then reconstruct expression profiles for genes along each of the edges using sliding windows summaries. Next we compute dot product scores for pairs of genes in edges (clusters) sampled at the same time or those representing the same pseudotime. Finally, we use permutation analysis to assign significance levels to the scores we computed. See Methods for details on each of the steps of TraSig.

**Figure 2:**
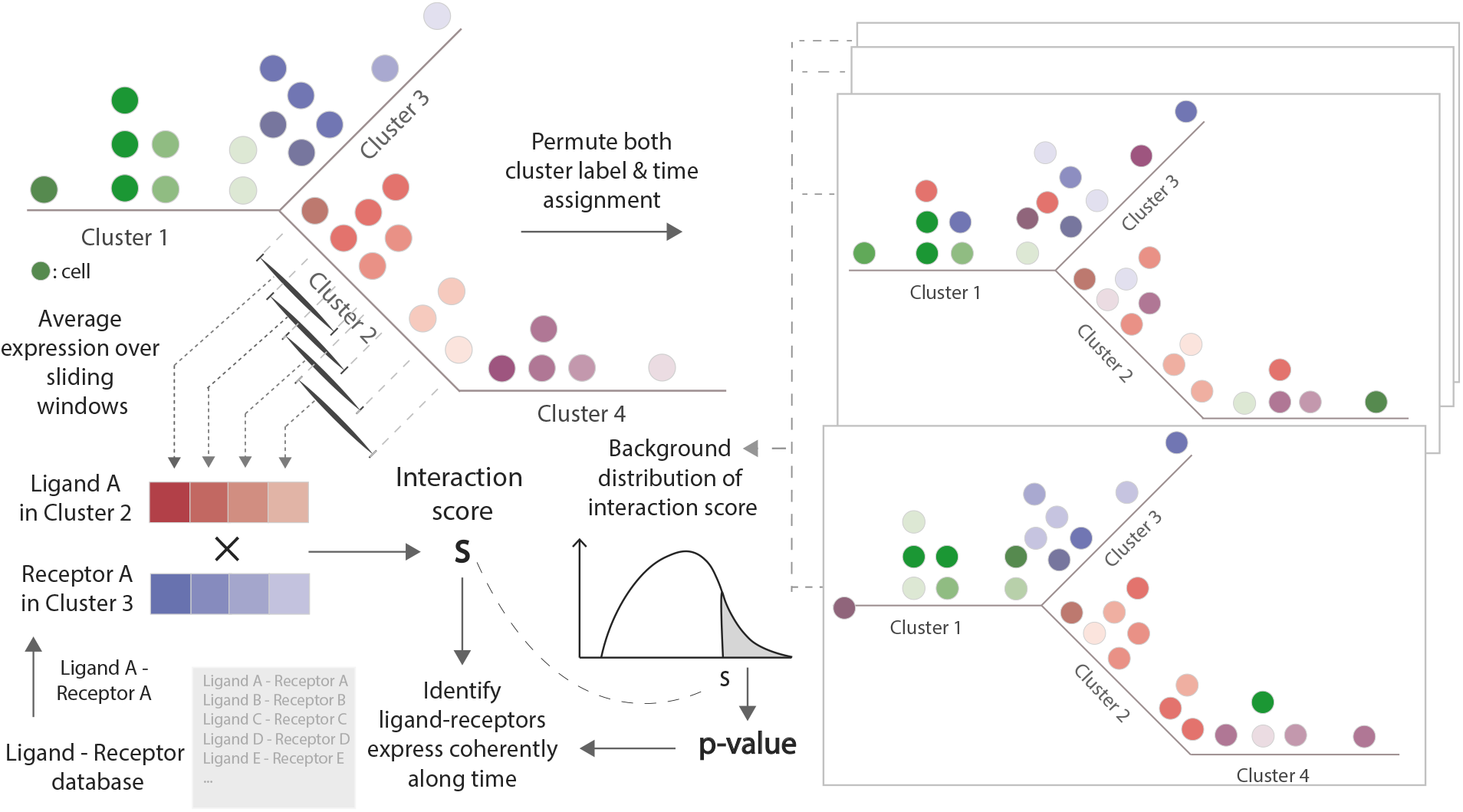
TraSig workflow. Top Left: For a scRNA-seq dataset, we use the reconstructed pseudotime trajectory and the expression data as inputs. Bottom Left: We next determine expression profiles for genes along each of the edges (clusters) using sliding windows and compute dot product scores for pairs of genes in edges. Right: Finally, we use permutation tests to assign significance levels to the scores we computed.

### Reconstructing dynamic liver development model using CSHMM

We first applied TraSig to a liver organoid differentiation scRNA-seq dataset composed of 11,083 cells sampled at two time points: day 11 and day 17 [11]. The data was preprocessed using a standard Seurat V3 [12] pipeline and cell types were assigned as previously discussed [11]. These were used to initialize trajectory inference using CSHMM [10]. Following filtering to remove genes not expressed in any of the cells, 26,955 genes were used to learn the CSHMM model. Figure 3a presents the resulting model learned for this data. As can be seen, the method identifies 12 clusters (edges) for these data. These agree very well with the clustering assignments from the Seurat single cell analysis. Specifically, CSHMM assigns separate edges for hepatocyte- (edge 3, 5, 9 and 10), endothelial- (edges 7 and 11), stellate- (edges 2 and 8), and ductal/cholangiocyte-like (edges 4 and 6) cells (Figure 4b). In addition, the model also presents informative pseudotime ordering of cells as we discuss below based on the reconstructed expression profiles for key marker genes.

**Figure 3:**
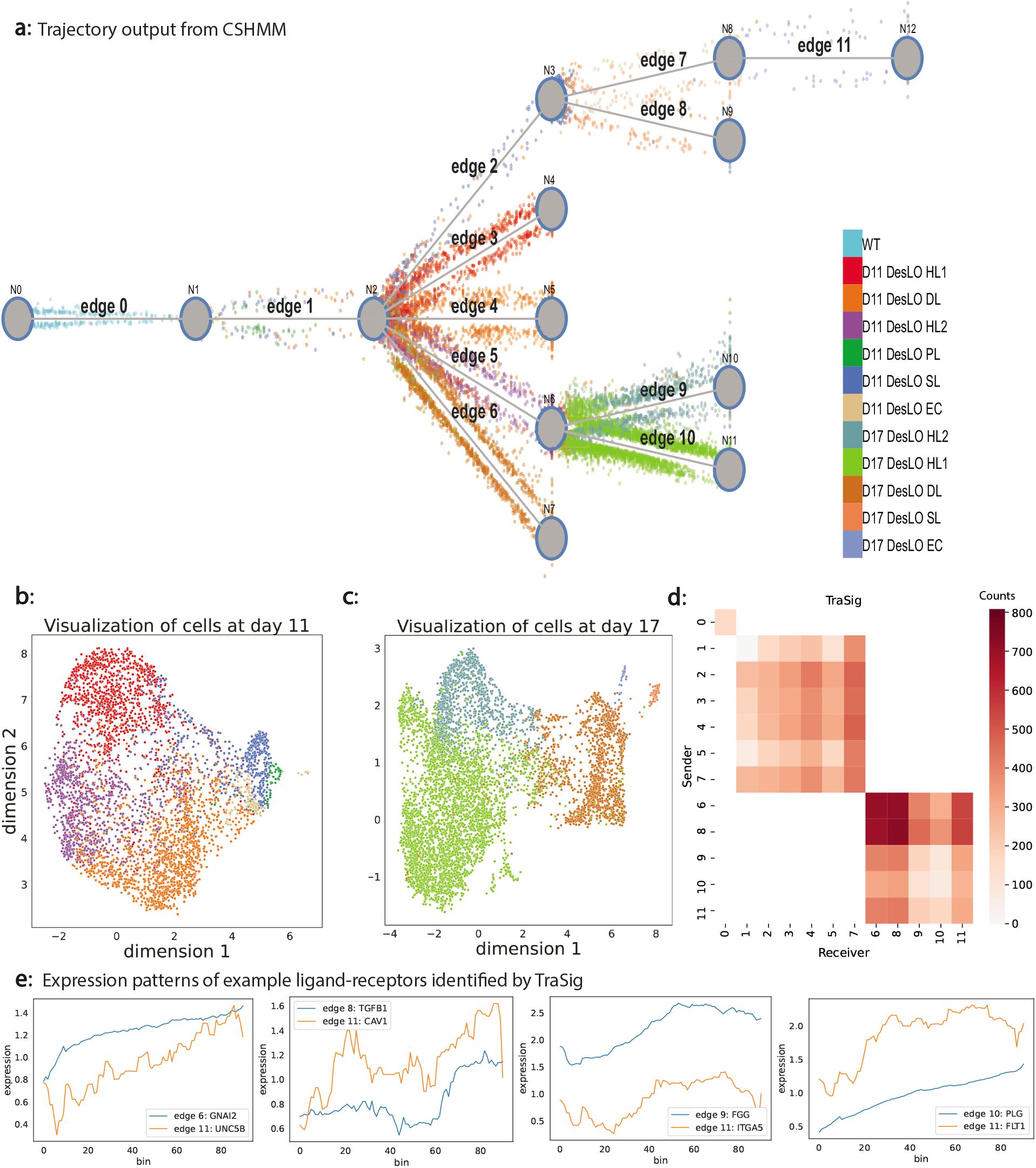
CSHMM and TraSig’s results on the liver organoid data (a) Reconstructed trajectory for liver organoid differentiation. CSHMM identifies a tree-structured trajectory that clusters cells to edges based on their expression pattern and relationship to the expression patterns of prior edges (Methods). Cells are colored by their cell type labels and are shown as dots ordered by their pseudo-time assignment. We also provide an interactive web user interface to better visualize the trajectory inference results (http://www.cs.cmu.edu/~trasig/). DesLO - designer liver organoid; HL - hepatocyte-like cells; DL - ductal/cholangiocyte-like cells; SL - stellate-like cells; EC - endothelial-like cells; PL – progenitor-like cells; WT - wild type. (b and c) UMAP [52] visualizations of the cells sampled at day 11 and day 17, colored by the cell type labels. (d) Heatmap for scores assigned by TraSig for all cluster pairs with cells sampled at the same time. (e) Sliding window expression for four example ligand-receptor pairs predicted to interact by TraSig.

**Figure 4:**
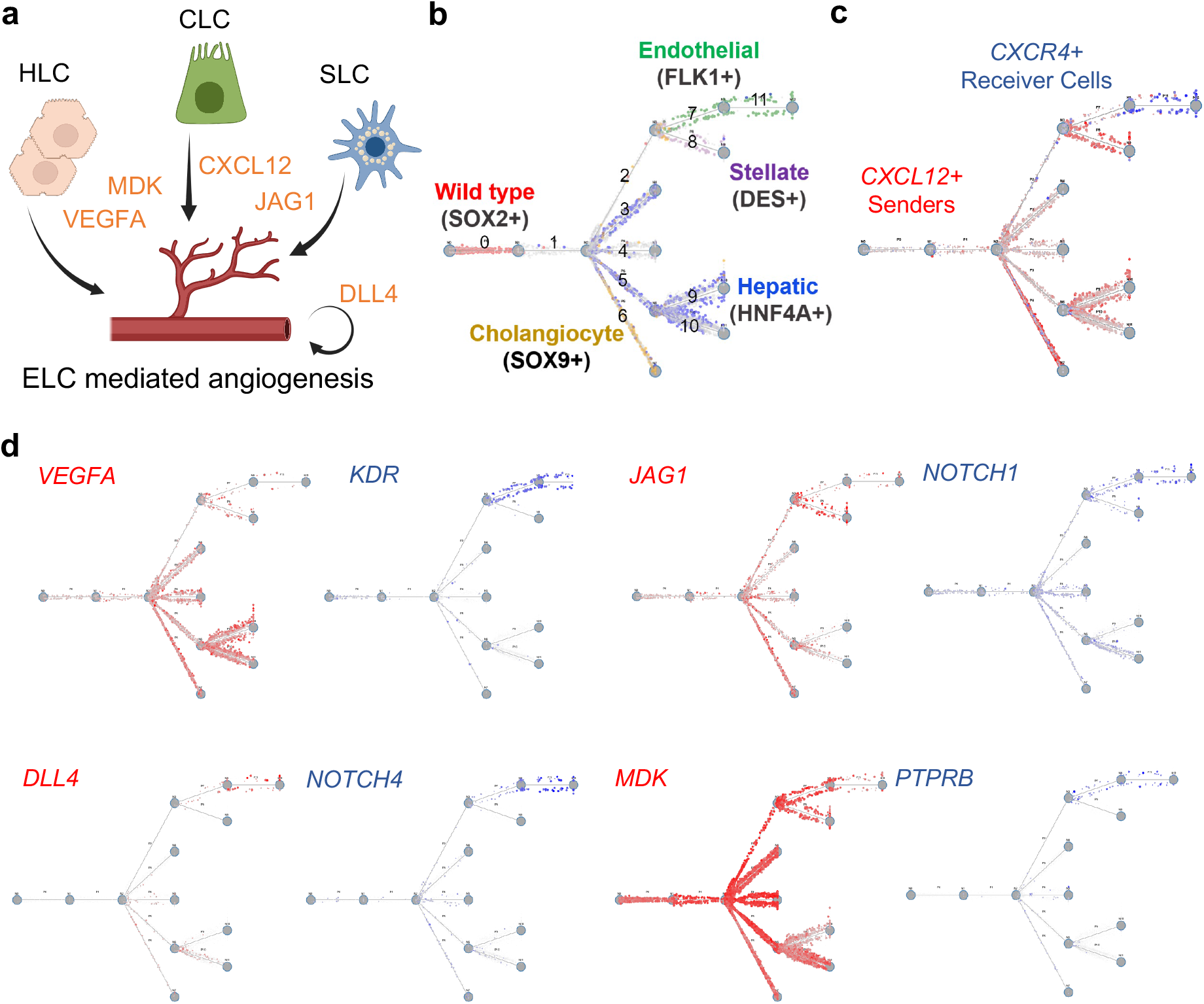
Ligand-receptor interaction predictions from TraSig of interest for functional studies. (a) Cartoon of cell signaling interaction between different DesLO cell types (HLC, hepatocyte-like cells; CLC, cholangiocyte-like cells; SLC, stellate-like cells; ELC, endothelial-like cells) (b) Trajectory plot showing cell type assignments with key identifying genes highlighted by different colors (Red = SOX2+ non induced cells, Yellow = SOX9 cholangiocyte-like cells, Blue = Hepatocyte-like cells, Purple = Stellate-like cells, Green = Endothelial-like cells). (c) Sender CXCL12 cells from the Cholangiocyte and Stellate populations in red shown with the receiver CXCR4 expressing endothelial cell population in blue. (d) Sender and receiver signaling populations (red = senders/ligands; blue = receivers/receptors). The darker the color is, the higher the expression level in a cell.

### Inferring cell type interactions for liver development

We next applied TraSig to the model reconstructed by CSHMM in order to gain insight into developmental signaling of co-differentiating liver cells from multiple germ layers. Such data is severely lacking for humans and so the use of the trajectory learned for liver organoid differentiation can provide valuable information on interactions regulating liver development. We thus tested all pairs of edges for which the assigned cells were from the same time point (Supplementary Notes). Figure 3d presents the results for scoring interactions between edges representing the same time (Methods). For the day 11 clusters (edge 1, 2, 3, 4, 5, 7), we find strong interactions between stellate-like 1 cells (edge 2) and endothelial-like cells (edge 7) and between ductal/cholangiocyte-like cells (edge 4) and endothelial-like cells (edge 7). For the day 17 clusters (edge 6, 8, 9, 10, 11), we find that the strongest interactions are between the ductal/cholangiocyte-like cells (edge 6) and stellate-like cells (edge 8). We also find high scoring interactions between stellate-like cells (edge 8) and endothelial-like cells (edge 11) and between ductal/cholangiocyte-like cells (edge 6) and endothelial-like cells (edge 11) for the day 17 clusters. The detection of significant interactions between the endothelial, stellate, and cholangiocyte cell types is further supported by their proximity in the liver. The stellate cells wrap around the endothelial cells and are bordered by the cholangiocyte comprised bile ducts [13].

### TraSig identifies ligand-receptor interactions important to vascular development

We evaluated the significant ligand-receptor pairs that were ranked highly by TraSig for the high scoring cluster pairs. We found that many agree with known functions and signaling pathways activated during liver development. Figure 3e presents a few examples of identified ligand-receptor pairs. We next studied the top scoring edges predicted to interact with endothelial-like cells. Endothelial cells play a major role in vascular development in liver [14]. To study the interactions of such cells, we looked for cluster pairs for which the receiver (receptor) cluster is the day 17 endothelial-like cell cluster (edge 11). GO term analysis of the identified ligands and receptors for these cluster pairs identifies several relevant functional terms related to vascular development including “blood vessel development” (minimum p-value among cluster pairs 5.72128*e* − 65), “regulation of endothelial cell proliferation” (p-value 3.34715*e* − 27) and “vascular process in circulatory system” (p-value 8.38655*e* − 12).

Many of the ligand-receptor pairs identified for interactions involving the endothelial-like cells are known to play a role in endothelial cell specification, migration, and angiogenesis further supporting the results of TraSig. Of note, we identified pairs including VEGFA/VEGFB/VEGFC with FLT1/KDR, which is required for proper liver zonation, sinusoid endothelial cell specification, and endothelial lipoprotein uptake [15, 16]; DLL4 with NOTCH1/NOTCH4, which is essential for endothelial tip and stalk cell crosstalk and liver sinusoidal endothelial cell capillarization [17, 18]; CXCL12 with CXCR4, which has been shown to promote endothelial cell migration and lumen formation independent of VEGF [19]; MDK with PTPRB, which is of great interest for its known impact on cancer angiogenesis [20, 21]; and CYR61 with ITGAV, which represents one of the many integrin interactions identified by TraSig which activate PI3K/AKT downstream signaling, and is known to regulate tip cell activity and angiogenesis (Figure 4a-d) [22].

### Experimental validation for predicted TraSig pairs

Given the success in identifying known interactions, we next experimentally validated additional TraSig predictions. We first assessed if there was a correlation between the signal level of CXCL12 or VEGF and vascularity via immunofluorescent staining of liver organoid cultures. As shows in Figure 5a-c, we found that loci with high relative expression of CXCL12 and VEGF co-localized with regions of increased vessel area percentage and vessel junction density, when compared to loci with relative low expression of CXCL12 and VEGF measured by AngioTool analysis of the immunofluorescent staining (see also Figures S5a and S5b).

**Figure 5:**
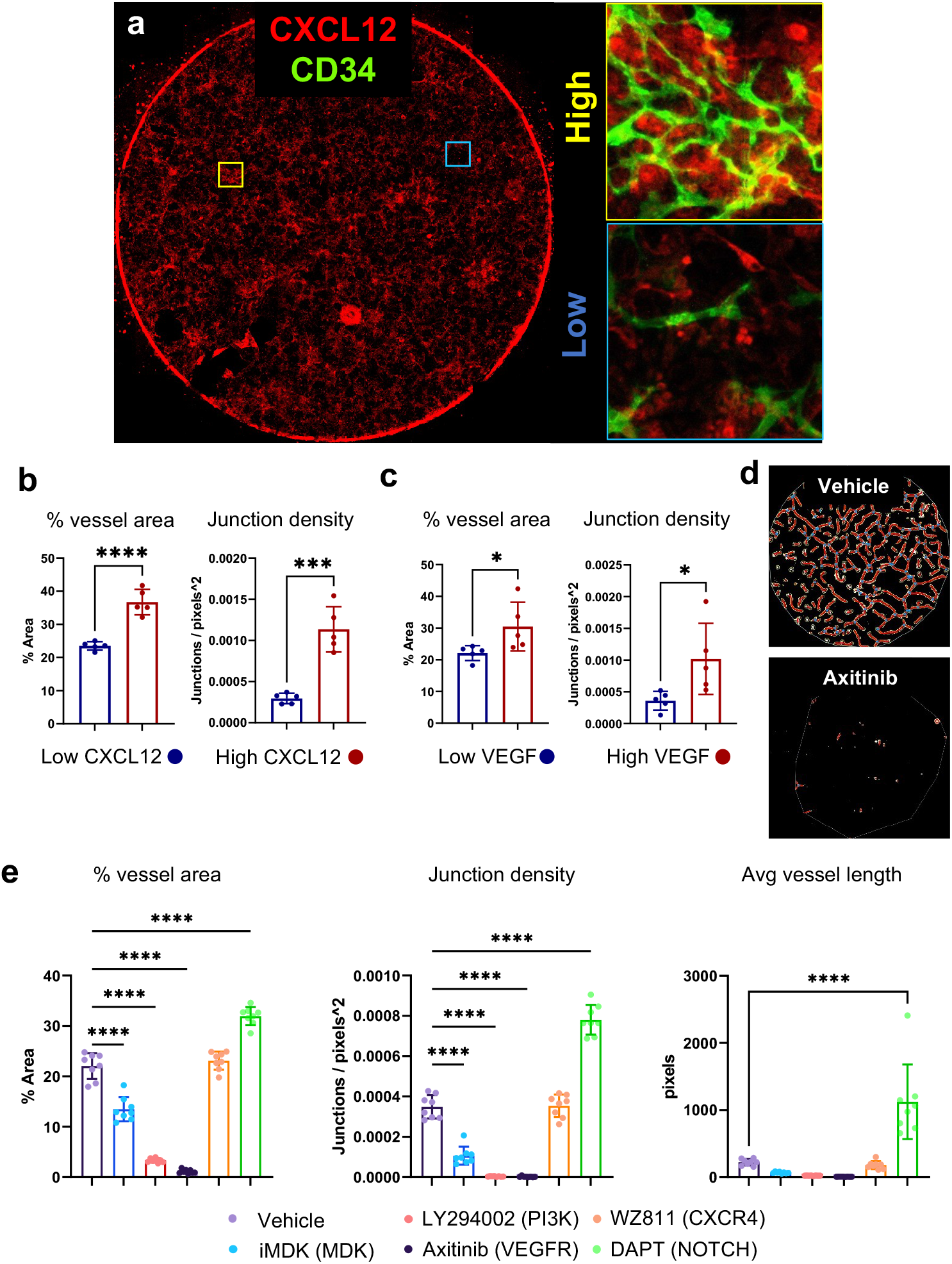
Functional validation of TraSig ligand-receptor signaling predictions. (a) Strategy for localized signaling effect of CXCL12. CXCL12 (red stain) overlaid with CD34 (green stain, on insets only) shown here with the yellow boxes indicating loci of high relative CXCL12 expression, and blue boxes indicating low relative CXCL12 expression. The same strategy was used for VEGF loci selection (see Methods). (b) Percent vessel area and junction density measured at CXCL12 and (c) VEGF low vs high loci from day 14 liver organoid cultures using AngioTool. n=4 loci for high CXCL12/VEGF expression and n=4 loci for low CXCL12/VEGF on one coverslip per staining combination. (d) Example of AngioTool evaluation of CD34 stained liver organoid cultures from the vehicle control (top) and Axitinib (bottom) conditions. (e) Percent vessel area, junction density, and average vessel length vascular metrics determined by AngioTool analysis results of CD34 stained liver organoid cultures with different inhibitor conditions. n=2 biological replicates with 4 sampled areas per coverslip. For b and c, Unpaired two tailed t test was used, * p<0.05, **** p<0.0001. For e, ANOVA with Tukey post comparison test was used, **** p<0.0001. Data are represented as mean ± SE for b, c, and e.

This motivated further investigation into the significance of predicted signaling interactions in the liver organoid cultures as they pertain to vascular development. We therefore performed prolonged (5 days from D9-14) inhibition of several predicted signaling proteins: VEGF, NOTCH, CXCR4, MDK, and PI3K (downstream of MDK and multiple integrin interactions). These experiments validated several of the predictions. Specifically, we observed significant decreases in percent vessel area, junction density, and average vessel length were detected in the VEGF, MDK, and PI3K conditions, while NOTCH inhibition revealed an opposite effect (Figure 5d and 5e). In contrast, the local correlation of increased vascular network formation with high CXCL12 expression did not carry over to a negative global effect via CXCR4 inhibition, indicating opportunity for further investigation, perhaps involving alternative inhibitors or assessment of the alternative CXCL12 receptor CXCR7, which also plays important roles in angiogenesis and liver regeneration [23, 24].

### Comparing TraSig with prior methods

We compared interactions predicted by TraSig to two popular methods for inferring cell type interactinons from scRNA-Seq data: CellPhoneDB [7] and SingleCellSignalR [8]. Both methods use the overall expression of genes in clusters and unlike TraSig do not use any ordering information. For both methods, we tested the same cluster pairs as we did for TraSig and used the same ligand-receptor database (Supplementary Notes). To make the comparisons more consistent, we combined the paracrine and autocrine predicted interactions for SingleCellSignalR since this is what other methods do. Figure 6a presents scores for all cluster pairs for TraSig, SingCellSignalR, and CellPhoneDB. As can be seen, while some pairs score high for all methods, others are only identified by one or two of the methods. Specifically, SingleCellSignalR seems to assign similar scores for most pairs whereas both TraSig and CellPhoneDB assign more variable scores. Figure 6c presents the Venn diagrams for the overlap between ligand-receptor pairs identified by the three methods for four example cell cluster pairs. In all cases, the receiver (receptor) cluster is the day 17 endothelial edge (edge 11). While SingleCellSignalR and TraSig overlap in roughly 50% of the identified ligand-receptor pairs, the overlap with CellPhoneDB is much lower.

**Figure 6:**
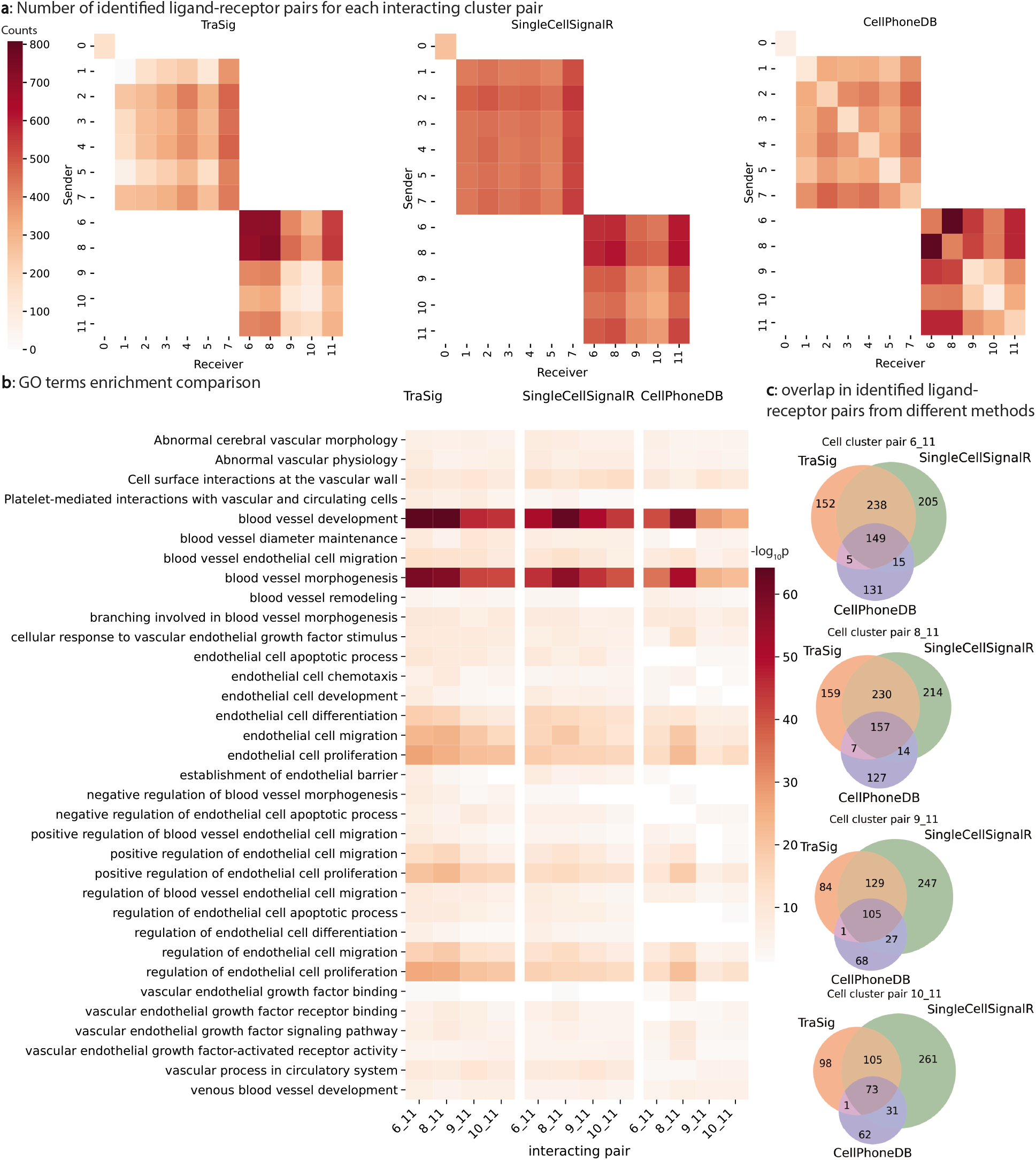
Results from comparing TraSig with SingleCellSignalR and CellPhoneDB. (a) Heatmaps for scores assigned by the three different methods for all cluster pairs representing cells sampled at the same time. (b) − log_10_ p-value for enriched GO terms related to endothelial cells and vascular development. (c) Venn diagrams for the overlap in identified ligand-receptor pairs among the three methods. The overlap between TraSig and SingleCellSignalR is high though roughly 50% of the identified pairs by each method are not identified by the other.

To evaluate the predicted pairs from these methods, we performed validation experiments, as mentioned above, and also compared enrichment p-values for relevant GO terms using ligands and receptors for several high scoring cluster pairs from each of the methods (See Supplementary Notes on how we select relevant GO terms). Among the significant ligand-receptors we successfully validated based on TraSig predictions, many were completely missed by CellPhoneDB even though they are included in the database it is using. These include DLL4-NOTCH1/4, JAG1-NOTCH1, VEGFB-FLT1 and VEGFC-KDR. As for SingleCellSignalR, for the DLL4-NOTCH1/4 predicted interaction SingleCellSignalR only identifies these as interactions within a single cell type and therefore does not identify the paracrine signaling between cell types. In contrast, TraSig identified these interactions as significant between day 17 endothelial-like cells (edge 11) and ductal/cholangiocyte-like cells (edge 6) and hepatocyte-like cells (edge 9 and 10). GO analysis further supports the advantages of TraSig. Figure 6b shows that TraSig leads to more significant relevant categories when compared to the two other methods. For example, TraSig obtains a minimum p-value among cluster pairs of 7.81570*e* − 60 for “blood vessel morphogenesis” whereas the minimum p-values for this category are higher for the other two methods (3.22968*e* − 57 and 6.02315*e* − 52 for SingleCellSignalR and CellPhoneDB respectfully). For “endothelial cell migration”, TraSig has a minimum p-value of 6.28812*e* − 25, again, lower than the minimum p-values for SingleCellSignalR (7.70322*e* − 20) and CellPhoneDB (2.06128*e* − 20). We obtained similar results when using another ligand-receptor database for all methods [25]. See Figure S13 for details.

### TraSig identifies interactions in neocortical development

To further evaluate TraSig’s performance, we applied TraSig to a mouse neocortical development scRNA-seq data [26]. After preprocessing (Supplementary Notes), we obtained 18,545 cells sampled at two time points: E14.5 and P0. We used the top 5000 dispersed genes to reconstruct CSHMM trajectories. The CSHMM model was initialized using the cell labels from [26]. Next the model was refined to improve both trajectory learning and cell assignment. The final trajectory learned for this data is presented in Figure S9. The model is composed of 44 clusters (edges) of which 23 contain cells from the first time point and 21 from the second. Next we applied TraSig to infer ligand-receptors pairs and interacting cluster pairs based on the sampling time.

Figure S7a presents scores for all cluster pairs. As can be seen, the method identified strongly interacting cluster pairs for both time points. The highest scoring interactions identified involve either endothelial cells (edge 18 from E14.5 and edge 39 from P0), radial glial cells (edge 1 from E14.5), interneurons (edge 24 from P0), or astrocytes (edge 26 from P0). We performed GO analysis using the significant ligands and receptors identified for radial glial cells in E14.5 or interneurons in P0. Figure S7b shows the – log_10_ p-value of enriched GO terms for interactions involving either RG2 [14-E] cluster for the radial glial cells in E14.5 (edge 1) or Int2 [14-P] cluster for the interneurons in P0 (edge 24). Radial glial cells were identified as progenitor cells for neocortical development [27] and determined to function as “scaffolds” for neuronal migration [28]. GO analysis shows that the signaling proteins identified by TraSig for interactions involving this cluster are indeed related to such functions and include “cell migration” (p-value 1.69780*e* − 60), “cell motility” (p-value 1.01291*e* − 56) and “regulation of cell migration” (p-value 9.23644*e* − 42). Terms related to neuron development are also highly enriched in the set of ligand and receptor proteins identified for the interneuron cell cluster and include “neurogenesis” (p-value 1.39908*e* − 64) and “neuron projection development” (p-value 5.39174*e* − 64).

### Applying TraSig to trajectories obtained by Slingshot

To test the ability of TraSig to generalize to pseudotime inferred by additional methods, we used it to post-process trajectories inferred by Slingshot [9]. Slingshot is a trajectory inference method that first infers a global lineage structure using a cluster-based minimum spanning tree (MST) and then infers the cell-level pseudotimes for each lineage. We applied Slingshot and TraSig to an oligodendrocyte differentiation dataset composed of 3,685 cells [29, 4]. Figure S8a presents the trajectory learned by Slingshot for this data. Figure S8b presents the interactions predicted by TraSig for the inferred trajectory. Cells assigned to edges 2 and 3 are more mature cells while those assigned to edges 0 and 1 containing precursor cells (Figure S8a). Our results suggest that the more mature oligodendrocytes are signaling to the precursors during development. As before we preformed GO analysis on the set of ligands and receptors predicted for strongly interacting clusters. We found several relevant GO terms including “neuron projection development” (p-value 2.50804*e* − 24) and “neuron development” (p-value 7.129894*e* − 23) (Figure S8c). Ligands in top ranking ligand-receptor TraSig pairs include PDGFA, BMP4 and PTN, all of which are know to be involved in regulating oligodendrocyte development [30, 31, 32].

## Discussion

Initial methods for the analysis of scRNA-Seq data mainly focused on within cluster or trajectory interactions. Recently, a number of methods have been developed to use these data to infer interactions between different cell types or clusters [6]. These methods focus on the average expression of ligands and their corresponding receptors in a pair of cell types to score and identify interacting cell types pairs.

While the exact way in which scores are computed differs between methods developed to predict such interactions, to date most methods looked at the average or sum of the expression values for ligands and receptors in the two clusters or cell types. Such analysis works well when studying processes that are in a steady state (for example, adult tissues) but may be less appropriate for dynamic processes. For real interactions, when time or pseudotime information is available, we expect to see not just average expression levels match but also trajectory matches in their expression profiles. Since many methods have been developed to infer pseudotime from scRNA-Seq data, such information is readily available for many studies.

To fully utilize information in scRNA-Seq data we developed TraSig, a new computational method for inferring signaling interactions. TraSig first orders cells along a trajectory and then extracts expression profiles for genes in different clusters using a sliding window approach. Matches between profiles for ligands and their corresponding receptors in different clusters are then scored and their significance is assessed using permutation tests. Finally, scores for individual pairs are combined to obtain a cluster interaction score. Since we use pseudotime ordering as input, we assume that the cells in the datasets we analyze are dynamically changing and that the input pseudotime ordering provides a good representation of the real time changes. We have experimentally tested that this is indeed the case for the liver organoid data we analyzed in this paper (Figure S11). We leave it up to users to decide if they would like to use the method for all cells profiled or for a subset of the cells (for example, those expected to change dynamically during the process being studied). Alternatively, we also provide an implementation of TraSig that following pseudotime ordering aligns the expression of cells in two edges (clusters) based on the expression of ligands and receptors. Next, the aligned profiles are used to score and identify interacting ligand-receptor and cluster pairs. See Supplementary Notes for details.

We applied TraSig to several different scRNA-Seq datasets and have also compared its predictions to predictions by prior methods developed for this task. As we have shown, for liver organoid development, TraSig was able to identify several known and novel interactions related to the regulation of vascular network formation. These interactions involve endothelial, stellate, and cholangiocyte cell types that have been known to reside in close proximity [13] and several ligand-receptor pairs known to be involved in vascular development. While many interactions were predicted by all methods we tested, there are also several interactions uniquely predicted by TraSig. We validated a number of these interactions including DLL4-NOTCH1/4, which are missed by CellPhoneDB and only identified by SingleCellSignalR as interactions within a single cell type. TraSig also uniquely identifies WNT2/3/4/7a/7b interactions with the FZD family and LRP6 supported by the known role of WNT in angiogenesis [33]. It also uniquely found BMP10-ACVRL1 / ACVR2A and SHH, interacting with multiple different receptors, both of which were also been shown related to angiogenesis [34, 35].

Our experiments showed that the VEGF inhibitor Axitinib, completely ablated the vascular network formation as shown previously [36, 11], and appeared to completely remove CD34 expressing cells. PI3K inhibition showed similar disruption of network formation, however, in contrast to Axitinib treatment, rounded CD34 expressing cells remained present and evenly spaced yet completely disconnected (Figure S5b). MDK inhibition appeared to decrease branching and connectivity of CD34 expressing cells significantly, however these cells still maintained a spread morphology. MDK is a pleiotropic growth factor that can induce cell proliferation, migration as well as angiogenesis [37, 38, 39]. It has been suggested that MDK from mesothelial cells can participate in liver organogenesis [40]. While its role was suggested in cancer related angiogenesis [41, 21], less is known about its function in liver development. Our combined computational and experimental analysis suggests such role for MDK in vascular development in human livers.

Interestingly, inhibition of NOTCH resulted in increased endothelial cell numbers and vascular formation. Vascularization can enable better engraftment in vivo. Hence modulation of notch signaling might be a possible target to improve liver organoid implantation in vivo that warrants further investigation. The mechanisms of these findings can be further investigated via cell type specific genetic circuits to determine dose, timing and cell types involved. Combined, our data confirms that significant signaling pathways in the liver organoids could be predicted using TraSig and functionally validated.

The INHBE-ENG interaction measured in the liver organoids (Figure S11b), was also found by TraSig. INHBE is uniquely highly expressed in primary liver as well as the liver organoids, and has been far less studied than it’s INHBA and INHBB counterparts [42]. Thus far, INHBE has been proposed as a hepatokine responsible for controlling energy homeostasis of white and brown adipose cells [43] and is potentially associated with insulin resistance [44], but has not been studied in the developing human liver to our knowledge. This poses a potential interesting avenue of further study that could help reveal the function of INHBE in liver, specifically as a regulator of angiogenesis during liver development.

Among the inhibitors we use, small molecules may have potential unintended off-target effects with limited spatial control. WZ811 and axitinib are relatively specific for inhibition of CXCR4 and VEGFR signaling respectively, while molecules like LY294002 can have broad effects due to the effects of PI3K signaling beyond its role downstream of integrin interactions. Likewise, DAPT, is a gamma secretase inhibitor that will prevent all NOTCH receptors from relaying downstream signals. Therefore, we view this as more of a proof of principle to test if TraSig is able to successfully determine natural key players important for angiogenesis in organoids.

We note that for this liver organoid data, the trajectory inferred by CSHMM put both edge 7 (mainly day 11 endothelial-like cells) and edge 8 (mainly day 17 stellate-like cells) downstream of edge 2, which mainly consists of day 11 stellate-like cells. This implicates the likelihood of common progenitor cells in edge 2, which can further differentiate into the endothelial lineage and pericyte(stellate) cells in liver organoids. In fact, co-development of pericytes in endothelial differentiation cultures has been observed recently [45], which may further suggest the presence of common mesodermal progenitors [46].

We have also tested TraSig on neuron and oligodendrocyte differentiation datasets. As we have shown, TraSig was able to correctly identify known and novel interacting cell types pairs for these datasets as well. For the first two datasets we studied, we used CSHMM for the pseudotime inference while for the oligodendrocytes, we applied TraSig to the pseudotime inferred by Slingshot [9]. This demonstrates the generalizability of TraSig which can be applied to output data from any pseudotime ordering method. As we have shown, the ability to identify significant interactions is independent of the ordering method itself enabling the use of TraSig in post-processing of any pseudotime ordered scRNA-Seq data.

## Methods

To identify interacting cell types pairs, we developed TraSig (**Tra**jectory based **Sig**naling genes inference), which infers key genes involved in cell-cell interactions. We primarily focus on genes encoding ligands and receptors at this stage but our method can accommodate other proteins likely to interact. For any two groups of cells that are expected to overlap in time, TraSig takes the pseudo-time ordering for each group and the expression of genes along the trajectory as input and then outputs an interaction score and p-value for each possible ligand-receptor pair.

### Learning trajectories for time series scRNA-Seq data

There have been several methods developed to infer trajectories from time series scRNA-Seq data [4]. Several of these methods first reduce the dimension of the data and then infer trajectory structures by using minimum spanning trees in the reduced dimension space [4]. While such methods work well for obtaining global ordering and for groupings cells, they may not be as accurate for the exact ordering of cells in the same edge (cluster), especially for clusters with small number of cells. Since the ordering is only based on the low dimension representation, genes that are only active in a small number of cells may have little impact on the representation of the cell in the lower dimension [10]. Since such ordering is critical for the ability to infer the activation or repression of individual genes along the pseudotime, we instead use another method for trajectory inference which works in the original gene space. This method, termed CSHMM, uses probabilistic graphical models to learn trajectories and to assign cells to specific points along the trajectories. CSHMM (Continuous-state Hidden Markov Model) [10] learns a generative model on the expression data using transition states and emission probabilities. CSHMM assumes a tree structure for the trajectory and assigns cells to specific locations on its edges. This enables both, the inference of the gene expression trajectories for each edge and the determination of overlapping edges (in time) which are potential interacting groups. In CSHMM, the expression of a gene *j* in cell *i* assigned to state *s_p,t_* is modeled as

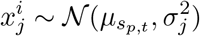
where *s_p,t_* is determined by both the edge *p* and the specific location *t* on the edge the cell is assigned to, and

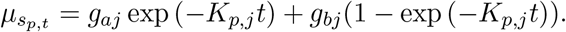

*g_aj_* and *g_bj_* are the mean expressions for gene *j* at branching node *a* and *b* (the beginning and the end of edge *p*, respectively) and *K_p,j_* is the rate of change for gene *j* on edge *p*. 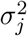 is the variance of gene *j*. CSHMM is learned by using an initial assignment based on clustering single cells and then iteratively refining the model and assignment using an EM algorithm [10].

### Selecting paired clusters

While most current methods look at all possible cluster pairs when searching for interactions, when using time series data we can constrain the search space and reduce false positives. Specifically, cells can only interact if both are active at the same time. For example, predicting interactions between clusters representing cells in day 1 and day 30 in a developmental study is unlikely to lead to real signaling interactions. TraSig can either use the time in which cells were profiled for this or it can use the tree structure provided by CSHMM to match edges based on their predicted pseudotime. Interactions are only predicted for pairs of edges (clusters) representing overlapping time.

### Ordering cells and inferring expression profiles

Given two groups of cells (cells assigned to two edges in the model) selected as discussed above, we first obtain a smooth expression profile for each gene along each of the edges. For this we first divide each edge into 101 equal size bins. We then use a sliding window approach that summarizes expression levels for genes along overlapping windows of equal size. We tested window sizes comprising of *L* = {5, 10, 20, and 30} bins and found that window size of 20 works best (Supplementary Notes). Windows overlap by *L* − 1 bins so the first *L* − 1 bins of a window are the last *L* − 1 bins of its predecessor. Since most cells are usually assigned to locations that are near the branching nodes (start and end of the edges, Figure 3a), we use *L*/2 as the length of the first sliding window and then increase to *L* when we reach the first *L* bins (Figure 2). We next generate an expression profile for each gene using its mean expression within each window. Using overlapping intervals allows us to overcome issues related to dropout and noise while still obtaining an accurate profile of the expression of the gene along the edge.

### Computing interaction scores for ligands and receptors

We used genes determined to be ligands or receptors from Ramilowski et al [47]. This database consists of 708 ligands, and 691 receptors with 2,557 known ligand-receptor interactions. To calculate an interaction score between a ligand in group A (sender) and its corresponding receptor in group B (receiver), we use the expression profile for each edge calculated as discussed above. Denote the expression values of the ligand in group A as x = (*x*_1_, *x*_2_,…, *x_M_*) and those for the receptor in group B as y = (*y*_1_, *y*_2_,…, *y_M_*), where *M* is the total number of overlapping intervals. We use the dot product function to compute the score by calculating 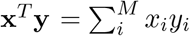. The advantage of using dot product for such analysis is that it enables the use of both the magnitude and the similarity of expression’s change over time to rank the top pairs.

To compute a p-value for the score, we use randomization analysis. Specifically, we permute the assignment of cells to edges and pseudotime in the model and re-compute the score as discussed above for the same pair of genes along the two clusters. Such permutation allows the method to identify interactions that are both, cluster (or cell type) specific and time dependent since genes that are active in most of the clusters will likely be also ranked high when permuting assignments between the clusters. We perform 100,000 permutations leading to a minimum p-value of 0.00001. We use Benjamini-Hochberg to control the false discovery rate (FDR) at 0.05 for multiple testing correction. For each pair of clusters, we also provide a summary score over all ligand-receptor pairs by counting how many ligand-receptor pairs are significant for this cluster pair.

### Alignment between paired clusters

The interaction score calculated as described above assumes that the cell clusters (edges) fully overlap in terms of their real time trajectory. While this assumption holds for many studies including for the data we analyze in this paper (Figure S11), there could be cases where the pseudotime represents different real time for different clusters or edges. To enable the use of TraSig in such cases, we also implemented another way of calculating the interaction score for TraSig. This option starts by obtaining the optimal *aligned* expression profiles for each pair of clusters (edges). By aligning clusters, we obtain the matching between the real time rather than the pseudotime dynamics of the two clusters. Next, we compute the dot product using the aligned profiles. The alignment method we used is adapted from those developed for bulk data [48, 49], based on B-spline interpolation and dynamic time warping (DTW). See Supplementary Notes for details.

### Using trajectories inferred by other methods

While we mainly discuss the use of TraSig with CSHMM, as we show in Results, it can be used with the output of any other trajectory inference tool. For this TraSig uses dynverse [4], which provides an R package that transforms the output of several popular trajectory inference and pseudotime ordering methods to a common output. Specifically, TraSig uses the “milestone_progression” output from dynverse which represents the location of a cell on an edge. This is a value in [0, 1] which we use to determine the pseudo-time assignment for each cell on an edge. All other steps are the same as when using CSHMM’s trajectory output. TraSig can also directly use pseudotime time and edge (cluster) assignment inputs from users if they prefer not to use the dynverse package.

### Assessment of cell-cell interaction to probe vascular formation in liver organoids

For evaluation of whole culture vascular network formation, liver organoids were cultured on 8 mm glass coverslips in a 48 well plate [11]. On day 9 of culture, indicated inhibitors 50 ng/mL VEGFR inhibitor, Axitinib (Sigma, Cat PZ0193-5MG); 15 uM CXCR4 inhibitor, WZ811 (Cayman, Cat 13639); 10 uM NOTCH inhibitor, DAPT (Stem Cell Technologies, Cat 082); 10 uM PI3K inhibitor, LY294002 (Stem Cell Technologies, Cat 72152); 1 uM MDK inhibitor, iMDK (Millipore, Cat 5.08052.0001); or vehicle control (DMSO, Sigma, Cat D2650-100mL) were supplemented to the culture medium daily for 5 days. After fixation with 4% PFA for 20 minutes at room temperature on day 14, the cultures were washed 3x in PBS and stained as explained previously [11] with CD34 antibody (Abcam, Cat ab81289) and the whole coverslip was imaged using an EVOS M7000. Raw images were exported to ImageJ and applied a threshold to generate binary images of the CD34+ vasculature networks. Four 1200 pixel ( 2-3 mm) diameter circular areas were selected per coverslip for assessment in AngioTool (https://ccrod.cancer.gov/confluence/display/ROB2) [50]. For evaluation of CXCL12 and VEGF localized vascular network formation, liver organoid cultures were fixed on day 14 and stained for CD34 along with either CXCL12 or VEGF. Loci, which we define here as 300 pixel diameter areas with high and low relative CXCL12 or VEGF expression determined by relative fluorescence, were identified in ImageJ and vascular network was analyzed using AngioTool.

## Availability of data and materials

TraSig is implemented in Python and is available at https://github.com/doraadong/TraSig. Single cell data for the liver organoid is available from the Gene Expression Omnibus (GEO) under accession number GSE159491. Single cell data for neocortical development [26] is available from the Gene Expression Omnibus (GEO) under accession number GSE123335. Single cell data for oligodendrocyte differentiation and for hepatoblast differentiation [29, 51, 4] are downloaded from https://doi.org/10.5281/zenodo.1443566.

## Acknowledgements

Figure 4a was created with Biorender.com. Figure S8a and Figure S10a were created using dynverse [4]. We also thank Haotian Teng for the fruitful discussion.

## Authors’ contributions

D.L., J.D., Z.B.-J. designed the research; D.L., J.D., Z.B.-J. developed the method; D.L. implemented the software; All authors analyzed the method outputs to select validation experiments. J.J.V., J.H. and M.R.E. designed and performed the validation experiments; D.L. and J.J.V. performed the analysis of validation data; All authors wrote the manuscript.

## Ethics approval and consent to participate

Human induced pluripotent stem cell work performed in this study were approved by the University of Pittsburgh Human Stem Cell Research Oversight (hSCRO) committee.

## Consent for publication

Not applicable.

