## Supplementary Notes and Figures for "Inferring cell-cell interactions from pseudotime ordering of scRNA-Seq data"

#### Data preprocessing for TraSig

For the liver data, we used Seurat [1] for data preprocessing. Specifically, we removed low quality cells by setting cutoffs for parameters "nFeature_RNA" and "percent.mt" and then normalized the counts using "LogNormalize" option (which composed of first calculating TPM with a scaling factor of 1e4 and then transforming by natural log).

For the neocortical development data, we used in-house normalization. We first filter out genes not expressed in any of the cells, then normalizing the counts using TPM with a scaling factor of 1e4 and finally transforming the values using natural log.

For the oligodendrocyte data and the hepatoblast data, we used the "expression" matrices of the datasets named "oligodendrocyte-differentiation-clusters_marques.rds" and "hepatoblast-differentiation_yang.rds" from dynverse [2] respectively and did not apply any extra preprocessing steps.

#### Data preprocessing for SingleCellSignalR and CellPhoneDB

For SingleCellSignalR, we used the same preprocessing procedures as for TraSig. For CellPhoneDB, given that the authors recommend using un-log transformed data [3], we applied the same preprocessing steps as for TraSig except for the last log transformation step.

#### Details in dividing edges into discrete bins

Given a continuous pseudo-time assignment in $[0, 1]$, we first round the original pseudo-time to 2 decimal places. This is equivalent to dividing each edge into 101 bins, where the middle 99 bins are of size 0.01 while the first and the last bins are of size 0.005.

#### Selecting the best window size for smoothing expressions profiles

We evaluated several options for the size of the sliding window. We compared windows of size: 5, 10, 20, and 30 bins. As expected (Figure S1), the larger the window, the smoother the resulting expression profiles. However, as the left-most plot of Figure S2 shows, the number of significant ligand-receptor pairs can decrease as the window size increases though this may be due to the permutation test which would be less sensitive to individual values when window size is larger. We thus looked for a size in which the number of significant pairs identified stabilizes. The middle and the right-most plots in Figure S2 present comparisons between the intersection ratio (y axes value) of significant ligand-receptors for different window sizes (x axis). As can be seen, once we reach 20, the number seems to stabilize and results are largely similar between 20 with both smaller and larger sizes.

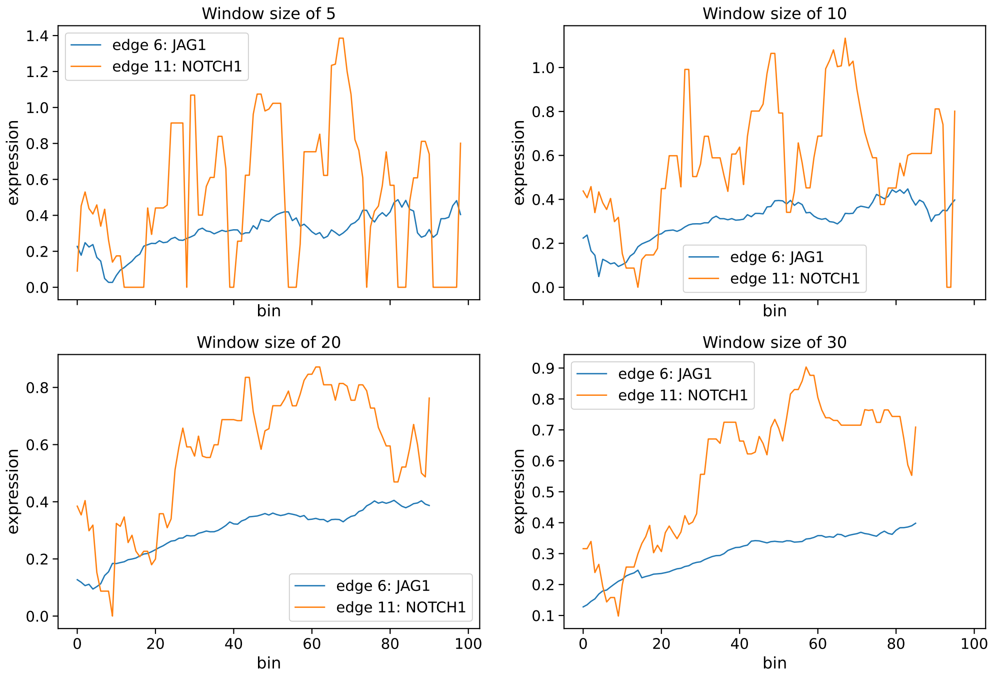


Figure S1: Expression profiles of an example pair of ligand-receptor under different window sizes for the liver organoids data. As the window size increases, the expression profiles become smoother. Note that the total number of sliding window intervals are different under different window sizes. Please refer to section "Details in calculating sliding window summaries" for details.

#### Details in calculating sliding window summaries

We implemented different options for sliding window summaries. Each of them differs in the way dealing with the first few sliding window intervals. When using a window of length $L$ to slide over $N$ bins, the first sliding window summary is taken over the first bin till the $L$th bin. Thus, the total number of sliding window summaries of size $L$ is $N-(L-1)$. One may just take these summaries as the smooth express profile (corresponding to the "discard" option in our implementation). However, this way of implementation may not be suitable for the cases where most of the information is allocated near the ends. Because in the common way of implementation as described above, the

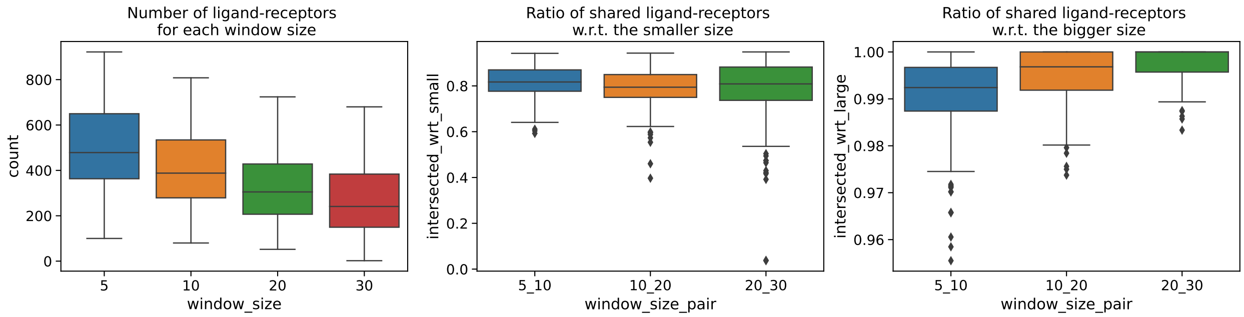


Figure S2: Impact of sliding window size on identified ligand-receptor pairs. Left: Number of significant ligand-receptors pairs for interacting clusters for different window sizes. Middle: Percent of significant ligand-receptors found by both window sizes in a pair (x axis) with regard to (w.r.t) all ligand-receptors found by the smaller window size. Right: w.r.t those found by the larger window size. The box-plots are generated using the standard definition (center line - median; box limits - lower and upper quartiles; whiskers - 1.5× interquartile range; points - outliers).

values at the ends (that is, cells closed to nodes in our case) are used fewer times than the values in the middle. In the output from trajectory inference tools, however, much more cells are assigned to locations closed to nodes (especially the starting node) than those in the middle.

To make better use of the values closed to the nodes and to obtain a higher resolution summary of the values around the starting nodes, we also implemented other options for taking sliding window summaries. The default option is "smallerWindow", where as described in the Methods, we use $L/2$ as the length of the first sliding window and then increase to $L$ when we reach the first $L$ bins. Another option is "None", where we use window size of 1 for the first bin, window size of 2 for the first 2 bins and gradually increase to window size of $L$ for the first $L$ bins. The last option is "parent", where we make use of the parenting edge of the edge of interest, and taking average by also using the cells assigned to the parenting edge. For example, when taking the summary for the first bin, we also use the cells assigned to the last $L-1$ bins in the parenting edge.

We tested different options of sliding window summaries on the liver organoid data and found they gave overall very similar predictions.

#### Calculating interaction scores using optimally aligned pseudotime expression profiles

Other than calculating scores directly using end-to-end alignment of the pseudotime expression profiles as described in the Methods section, we implemented another option by first obtaining the optimal temporal alignment and then calculate the scores based on the aligned profiles, adapted from [4, 5]. The main steps for the alignment are shown in Figure S3.

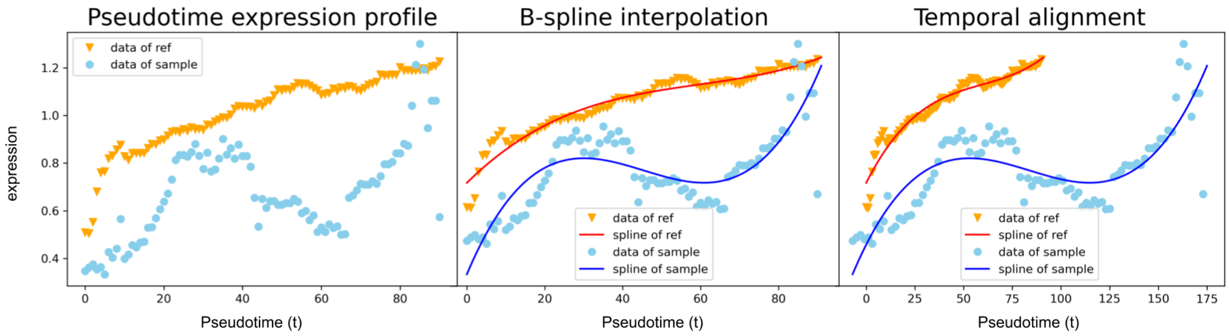


Figure S3: Obtain the optimal alignment for every pair of edges / clusters. One of the edge / cluster is called the reference (ref in the figure) and the other is called the sample. Here we illustrate the main idea using a single pair of genes. We start with the sliding window summaries of these genes along pseudotime (left). We first find the B-spline representations of these expresssion profiles (middle). We finally obtain the optimal alignment between these two edges / cluster by minimizing the differences between the aligned profiles (right).

For every ligand and receptor in the database and for every edge / cluster, we first find the B-spline representation of the sliding window summaries. We use the interpolate.splrep function from scipy package [6], with degrees set as 3 (cubic splines) and the smooth parameter as 10 for the liver organoid data.

We use dynamic time warping (DTW) to align every pair of edges / clusters for which we will calculate interactions scores. For each pair of edges / clusters, we use the sender edge / cluster as reference and align it with the receiver edge / cluster. Specifically, we assume a linear alignment function for each pair $\tau_j(t) = \frac{(t-b_j)}{a_j}$, following [4, 5], where $t$ is the time in the reference, $j$ is the $j$th pair of edges / clusters and $a_j$ and $b_j$ are parameters for this function. The objective of this alignment step is to find the best $a_j$ and $b_j$ for the $j$th pair of edges / clusters. Particularly, we find them by looking at the differences between the aligned expression profiles of the reference (sender) and the receiver edge / cluster and select the set of $a_j$ and $b_j$ minimizing the differences.

We focus on the expression differences between ligands in the reference (sender) edge / cluster and their corresponding receptors in the receiver edge / cluster. For the $j$th pair of edges / clusters and for the $k$th pair of ligand and receptor, we first obtain their aligned expression profiles. Set $s^j_{l_k}(t)$ as the expression of the ligand $l_k$ at time $t$, estimated from the corresponding fitted spline.

Here $t \in [t_{min}, t_{max}]$, the reference interval, and we set $t_{min}$ as 0 and $t_{max}$ as the total length of the sliding window summaries. Similarly, the expression of the ligand's receptor $r_k$ at $t'$ is $s^j_{r_k}(t')$, where $t' \in [t_{min}, t_{max}]$. Then the aligned expression for $r_k$ is $s^j_{r_k}(\tau_j(t))$. The aligned interval is thus $[\alpha, \beta]$, where $\alpha = \max(t_{min}, \tau_j^{-1}(t_{min}))$ and $\beta = \min(t_{max}, \tau_j^{-1}(t_{max}))$. Given the fitted splines, we can estimate the expression values at any $t \in [\alpha, \beta]$. We use cosine distance defined as $1 - \frac{\mathbf{l}^T\mathbf{r}}{||\mathbf{l}||||\mathbf{r}||}$ to measure the expression differences, where $\mathbf{l} = \{s^j_{l_k}(\alpha), ..., s^j_{l_k}(t), ..., s^j_{l_k}(\beta)\}$ and $\mathbf{r} = \{s^j_{r_k}(\tau_j(\alpha)), ..., s^j_{r_k}(\tau_j(t)), ..., s^j_{r_k}(\tau_j(\beta))\}$. The overall differences between all pairs of ligands and receptors for the $j$th pair of edges / clusters is just the summation over all pairwise distances.

To avoid trivial solutions, we set the following constraints: $\alpha > 0$, $\alpha < \beta$ and $\frac{(\beta-\alpha)}{(t_{max}-t_{min})} \geq \epsilon$. The last constraint keeps the overlap between the aligned interval and the reference interval at least $\epsilon$. We use $\epsilon = 0.5$ for the liver organoid data.

For the $j$th pair of edges / clusters, after we find the optimal alignment function (the function having the best $a_j$ and $b_j$ values), we then calculate the interaction score for each ligand-receptor pair using the expression profiles aligned under this function. When estimate the significance level using permutation test, we apply the same optimal alignment function to the permuted samples and calculate interaction scores using the aligned expression profiles.

#### Assigning sampling time and cell type for clusters output from pseudotime inference tools

For a cluster / edge, we assign the cell type and sampling time based on the cell type labels and sampling time assigned to the majority of the cells in the cluster. We applied this to all datasets except for the cell type labeling of the oligodendrocyte data and hepatoblast data, where we also note down the cell type labels of all smaller groups that altogether make to 90% of the whole cluster / edge, other than the majority group.

#### Coloring cells in trajectory plots by expression values

The plots in Figure 4b-d were generated by coloring the cells according to their expression levels of a certain gene. The darker the color is, the higher the expression level in a cell. For all genes shown in Figure 4c-d, we set an expression cutoff of 1. The cell is shown as a bigger size if having the expression above 1 or a smaller size if not expressing the gene at all. The cell size is in the middle if it expresses the gene but the level is below 1. In addition, the opacity is set to 20% for cells not expressing the gene.

#### Use customized databases for SingleCellSignalR and CellPhoneDB

Given we want to use the same database for the comparison among methods and this database is different from the default databases used by SingleCellSignalR and CellPhoneDB, we need to construct customized databases for these two methods. Since the default database for SingleCellSignalR is also based on gene symbols, we just replace the default list of ligand-receptor pairs with ours. This is less straightforward for CellPhoneDB whose inferences are based on protein-protein interactions. We follow instructions provided by [3] to prepare this database. Particularly, we first prepare the "gene_input" by mapping all genes in our database to their UniProt IDs using the conversion lists provided by [7]. A few genes don't have corresponding UniProt IDs and are thus removed. For those genes mapped to multiple UniProt IDs, we keep only one single isoform to avoid duplicates. We then prepare "protein_input" and "interaction_input" accordingly.

We follow the same steps when we prepare the customized databases for the other database [7] we use.

#### Values used to anchor the colormaps for the heatmaps for methods comparison

To enable the comparison between different methods' output, we set the lower anchor value as the minimum interaction score (the least number of identified ligand-receptor pairs for a edges / clusters pair) across all three methods and the higher anchor value as the maximum interaction score (the largest number of identified ligand-receptor pairs for a edges / clusters pair) across all three methods. Specifically, the anchor values are set as 0 and 808 for the heatmaps shown in Figure 6 and 0 and 601 for the heatmaps shown in Figure S13.

#### GO analysis

We use both ligands and receptors of the identified ligand-receptor pairs as the input gene sets and use gProfiler for the GO term enrichment analysis [8]. Among all significant GO terms from all cluster pairs and comparison methods, we select the terms relevant to the biology process of interest for visualization. For the analysis on the liver organoid development data, we select the terms containing keywords “endothelial”, “vessel” or “vascular”. For the analysis on the neocortical development data, we select the terms containing keywords “Axon”, “Nervous”, “Neuroactive”, “axon”, “axonal”, “brain”, “nervous”, “neuron”, “neuronal”, “synapse”, “Notch”, “calcium”, “fibroblast”, “glial”, “ion”, “localization”, “migration”’, “motility”, “neurogenesis”, “neurotrophic”, “pluripotency”, “potentiation”, “stem”, “synaptic”, “transmission”. For the analysis on the oligodendrocytes data, we select the terms containing keywords “Development”, “development”, “developmental”, “differentiation”, “formation”, “localization”, “locomotion”, “migration”, “morphogenesis”, “motility”, “movement”, “multicellular”, “regeneration”. For the analysis on the hepatoblast data, we select the terms containing keywords “development”, “developmental”, “differentiation”, “formation”, “localization”, “locomotion”, “migration”, “morphogenesis”, “motility”, “movement”, “multicellular”, “vascular”, “vessel” and “neurogenesis”. When too many GO terms selected, we applied extra filtering steps. For the liver organoid data, we use the set of GO terms with minimum p-value smaller than $1.20508e-06$ in at least one of the three comparison methods for visualization. For the neocortical development, we list the set of GO terms with minimum p-value smaller than $5.35648e-40$ in at least one of the two clusters. For the oligodendrocytes data, we list the set of GO terms with minimum p-value smaller than $2.15260e-12$ in at least one of the two clusters. For the hepatoblast data, we list the set of GO terms with minimum p-value smaller than $6.14735e-15$ in at least one of the two clusters. For the analysis on the liver organoid data using the other database [7], we use the set of GO terms with minimum p-value smaller than $3.97532e-06$ in at least one of the three comparison methods for visualization.

Unless specified, the p-values listed in the parentheses for the GO terms are the minimum p-values among all interacting cluster pairs .

#### qPCR timelapse experiments for the validation of the assumptions of TraSig for liver organoids

Human liver organoids were generated as previously described [9] in 48 well plates. The tissues were harvested and lysed on days 7, 8, 9, 10, and 12 by adding $300\mu L$ Trizol (Invitrogen, Cat# 15596018) directly to the tissue culture well, pipetting several times, collecting, and storing at -80°C to be thawed when ready for extraction. RNA was extracted using the Direct-zol RNA miniprep kit (Zymo, Cat# R2052) according to manufacturer’s instructions. cDNA was synthesized using the high Capacity cDNA reverse transcription (Applied Biosystems, Cat# 4368813). qRT-PCR was performed using the PowerUp™ SYBR™ Green Master Mix intercalating dye (Applied Biosystems™, Cat# A25742) according to the manufacturer's instructions with 20ng total cDNA. Expression was normalized to 18S ribosomal RNA and relative gene expression was calculated using $2^{-\Delta\Delta CT}$ method. Primers used for qRT-PCR are listed in Table S1.

### Supplementary Figures

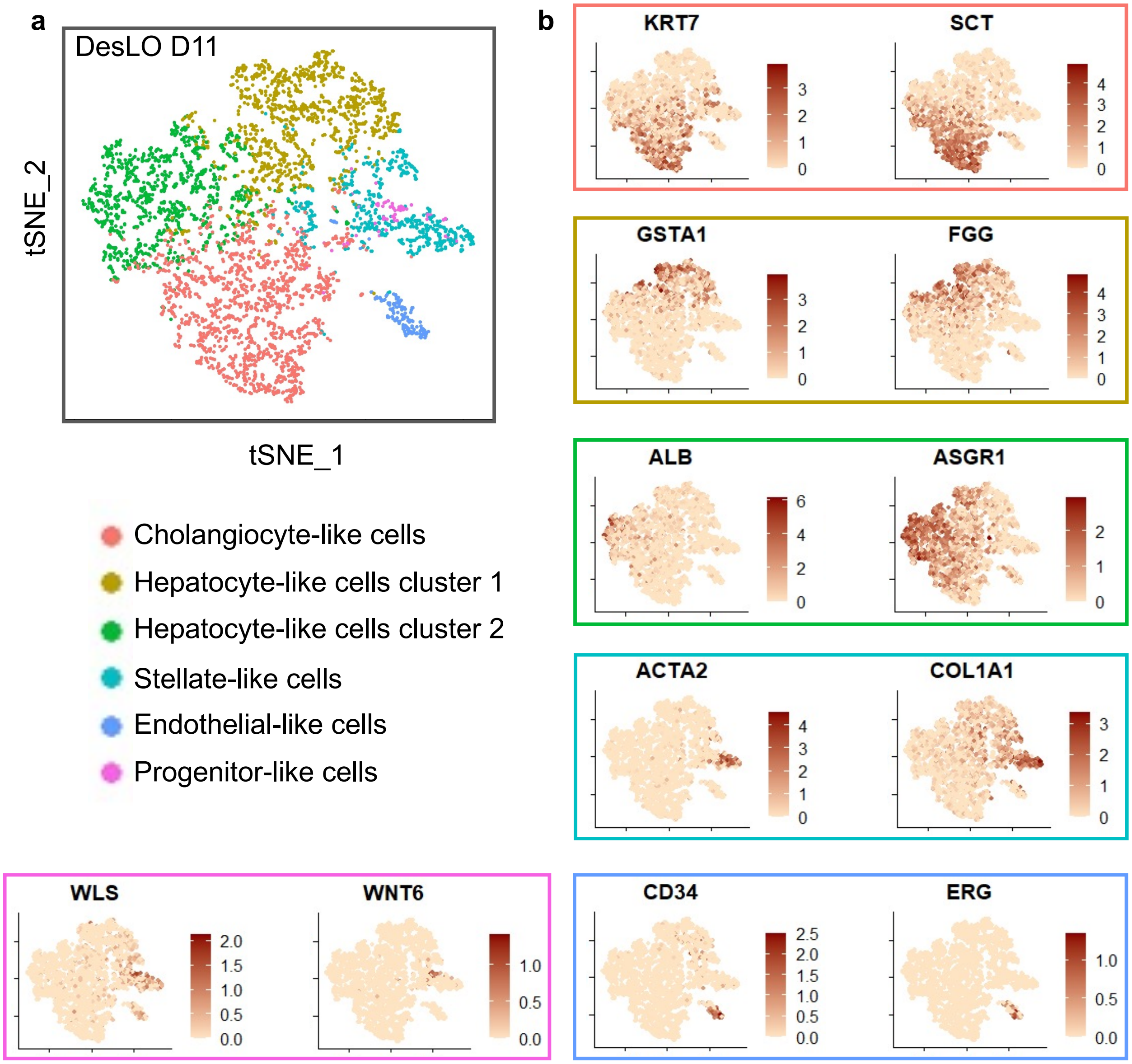


Figure S4: DesLO day 11 scRNA-Seq cluster analysis. (a) The tSNE plot showing clustering analysis of day 11 DesLO scRNA-seq sample revealed 6 distinct clusters, cholangiocyte-, stellate-, endothelial-, progenitor-, and two hepatocyte-like cell clusters based on differential gene expression. (b) Highly enriched genes of each cluster are indicated by colored boxes.

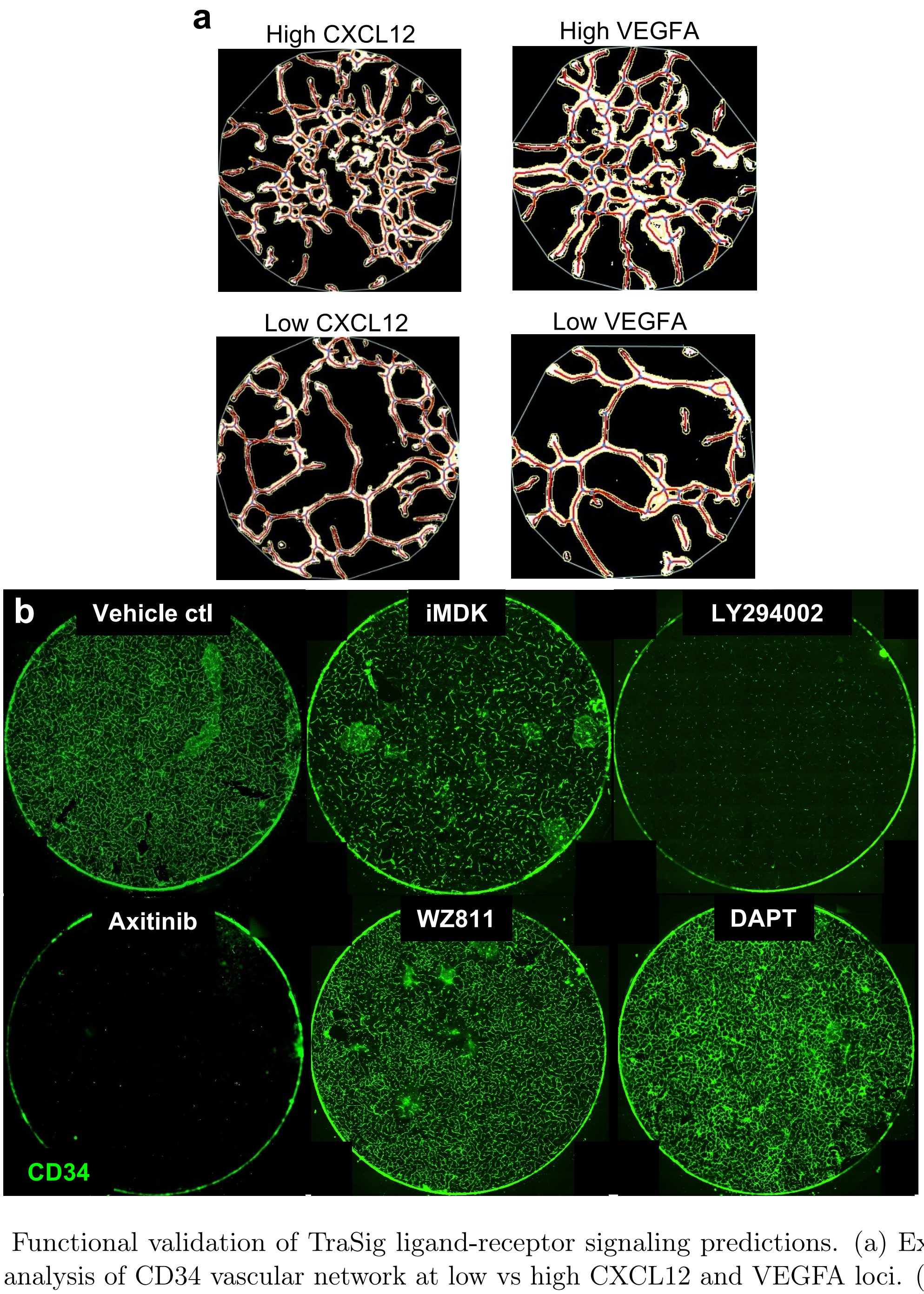


Figure S5: Functional validation of TraSig ligand-receptor signaling predictions. (a) Example of AngioTool analysis of CD34 vascular network at low vs high CXCL12 and VEGFA loci. (b) Whole culture image of 8 mm coverslip for 5 inhibition conditions implicated by TraSig ligand-receptor signaling predictions (representative images of n=2 biological replicates).

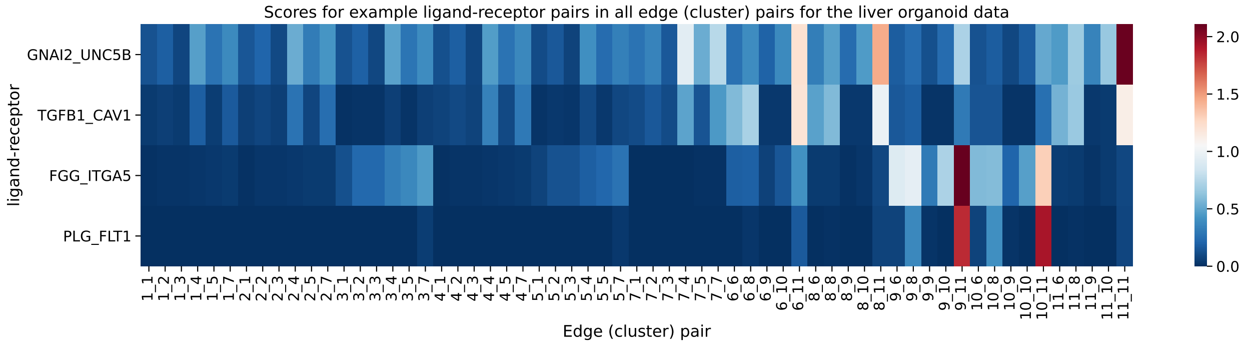


Figure S6: Scores in all edge (cluster) pairs for the example ligand-receptors shown in Figure 3e for the liver organoid data. These ligand-receptor pairs are significant for some edge (cluster) pairs and not others. The significant ones, for example, GNAI2-UNC5B in pair 6_11 and TGFB1-CAV1 in pair 8_11, are often scoring higher than many of the other edge (cluster) pairs.

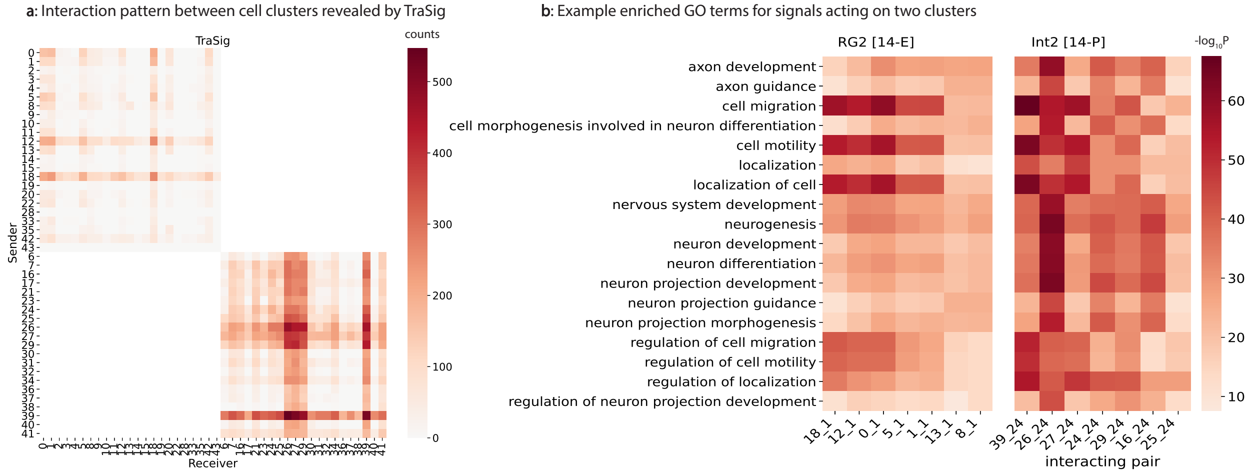


Figure S7: TraSig's results on the neocortical development data. (a) The strongest interactions identified are between endothelial cells (edge 18 in E14.5 (top) and edge 39 in P0 (bottom)) and other groups of cells including interneurons (edge 24 from P0) and radial glial cells (edge 1 from E14.5). (b) GO term enrichment analysis shows that TraSig identifies interactions specifically related to cell migration and neuron development.

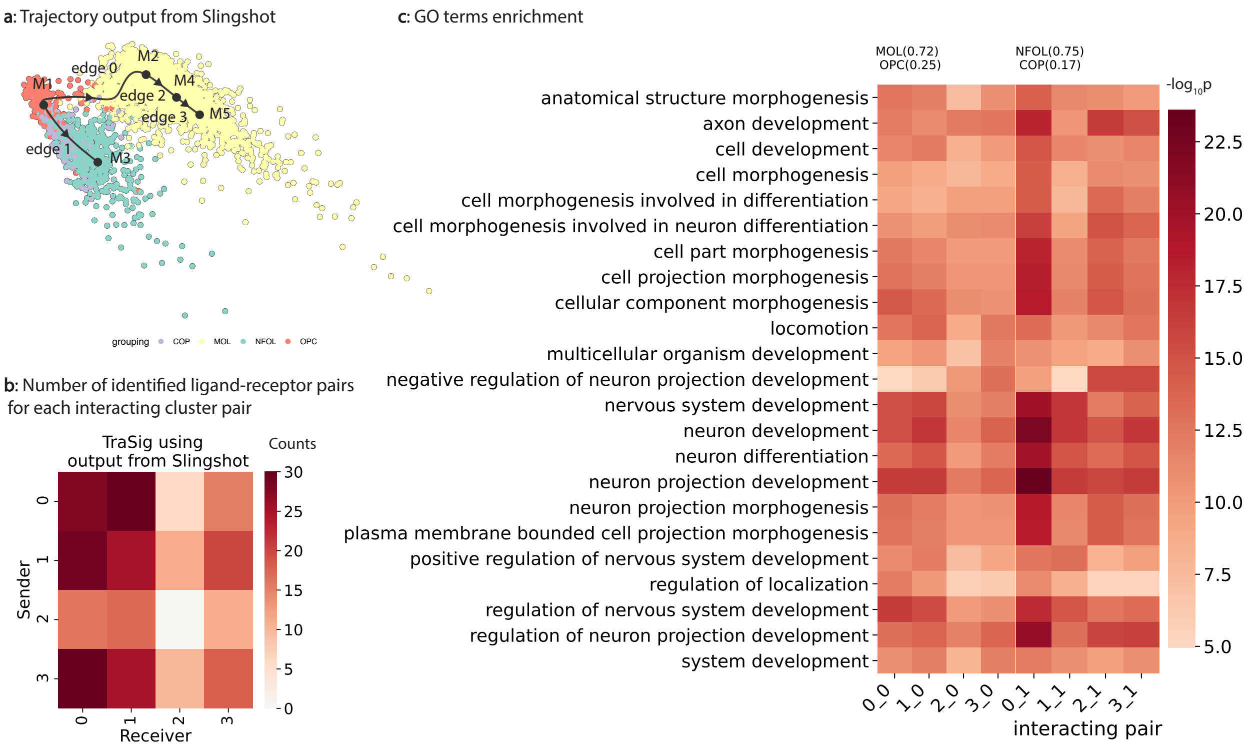


Figure S8: Application of Slingshot and TraSig to the oligodendrocytes development data. (a) Slingshot pseudotime ordering and trajectory inference. Cells are colored by the labels assigned to them in the original paper [10]. OPC - Pdgfra+ oligodendrocyte precursors; COP - Differentiation-committed oligodendrocyte precursors; NFOL - Newly-formed oligodendrocytes; MOL - Mature oligodendrocytes; the milestones (nodes) output from Slingshot are denoted by $M1, M2, ...$; we used dynverse [2] to generate the plot. (b) TraSig interaction scores for edges (clusters) pairs identified by Slingshot. (c) Enriched GO terms and $-\log_{10}$ p-value for strongly interacting cluster pairs. The first 4 interactions all involve cluster 0, which consists of 72% of MOL cells, 25% of OPC cells and other minor cell types, as noted by the column name. The next 4 interactions involve cluster 1, composed of 75% NFOL, 17% of COP cells and other minor cell types.

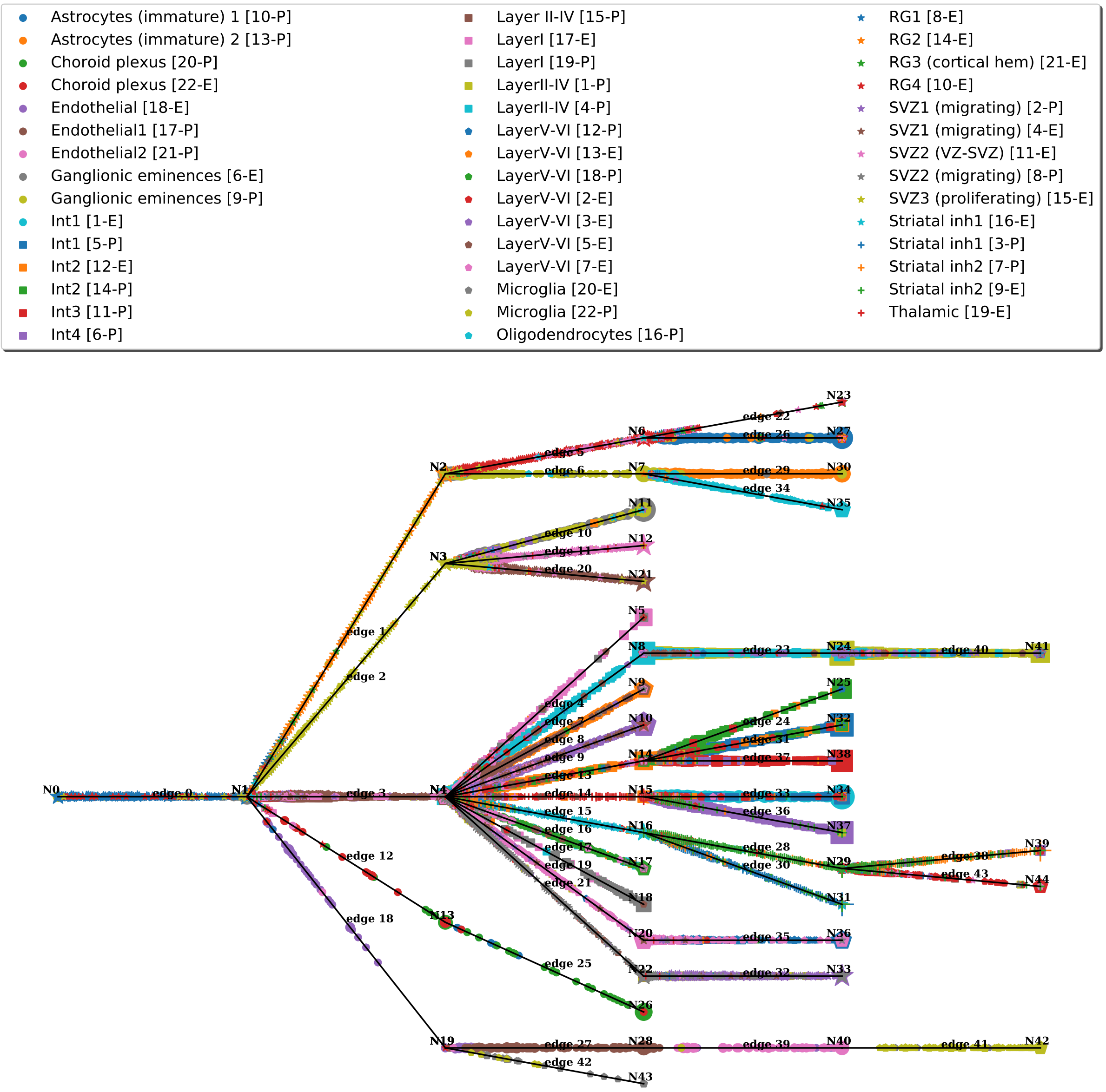


Figure S9: Reconstructed trajectory for neocortical development. CSHMM identifies a tree-structured trajectory that clusters cells to edges based on their expression pattern and relationship to the expression patterns of prior edges (Methods). Cells are colored by their cell type labels assigned in the original paper [11] and are shown as dots ordered by their pseudo-time assignment.

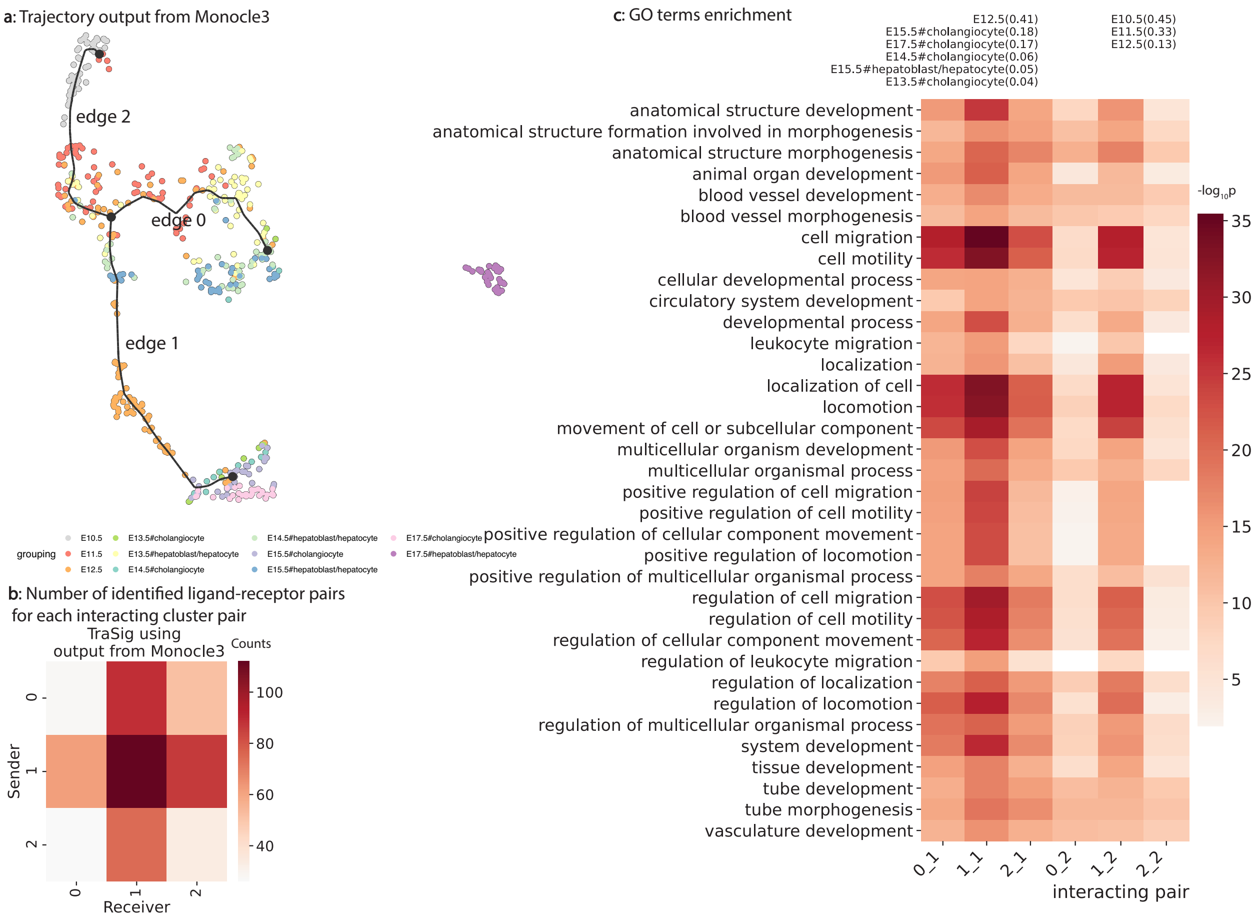


Figure S10: Application of Monocle3 and TraSig to the hepatoblast differentiation data [12]. (a) Monocle3 pseudotime ordering and trajectory inference. Cells are colored by the labels assigned to them in the original paper. We used dynverse [2] to generate the plot. (b) TraSig interaction scores for edges (clusters) pairs identified by Monocle3. (c) Enriched GO terms and $-\log_{10}$ p-value for strongly interacting cluster pairs. The first 3 interactions all involve cluster 1, which consists of 41% of E12.5 cells and 18% of E15.5#cholangiocyte cells and other minor cell types as noted by the column name. The next 3 interactions involve cluster 2, composed of 45% E10.5 cells, 33% of E11.5 cells and other minor cell types.

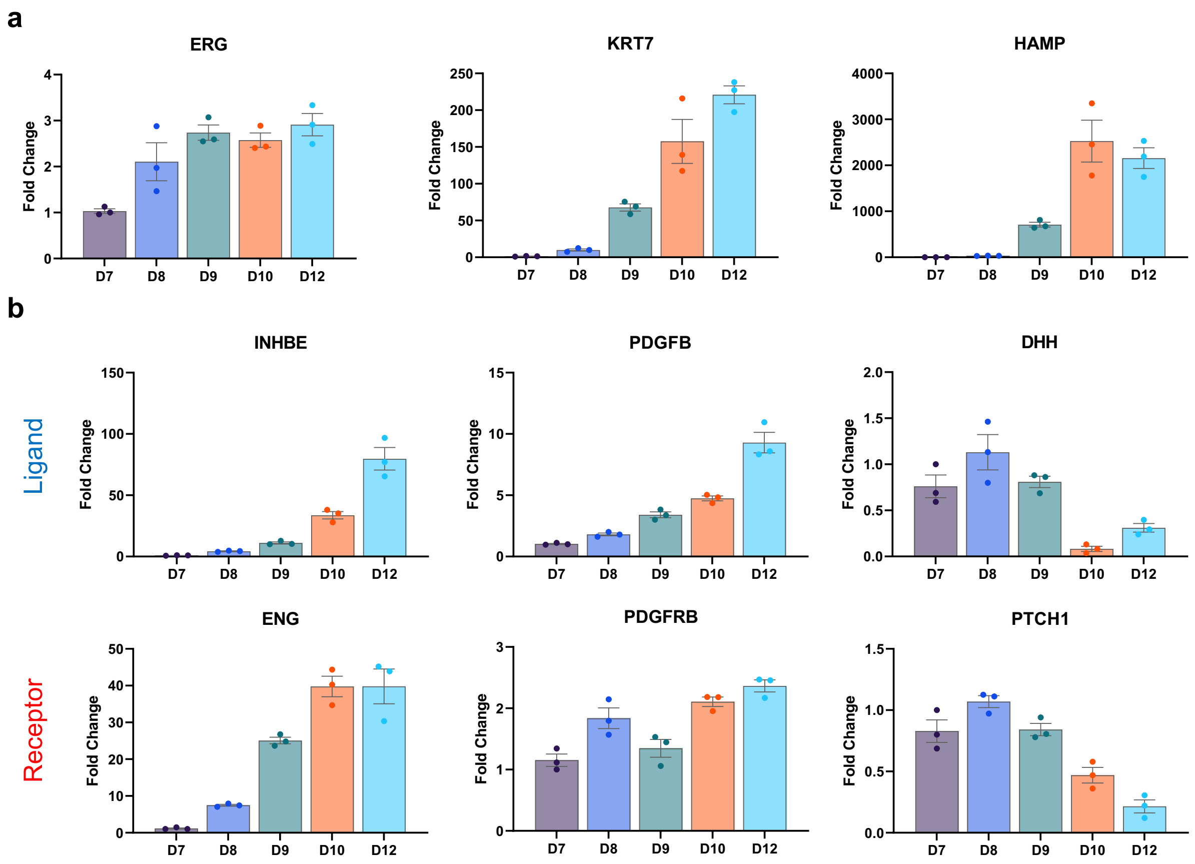


Figure S11: DesLO timelapse bulk RNA analysis of cell population markers and cell-cell signaling. DesLO samples were generated and their RNA were extracted at days 7, 8, 9, 10, and 12 of culture. (a) shows the qPCR determined fold-change (over day 7) gene expression of endothelial marker, ERG; cholangiocyte marker, KRT7; and hepatocyte marker, HAMP. The timelapse data shows relatively stable expression of ERG, while the aligned temporal emergence of cholangiocyte- and hepatocyte-like cells (indicated by expression of KRT7 and HAMP, respectively) appear on day 9 of culture. (b) displays the qPCR fold-change (over day 7) results for signaling pairs INHBE-ENG, PDGFB-PDGFRB, and DHH-PTCH1 identified by TraSig.

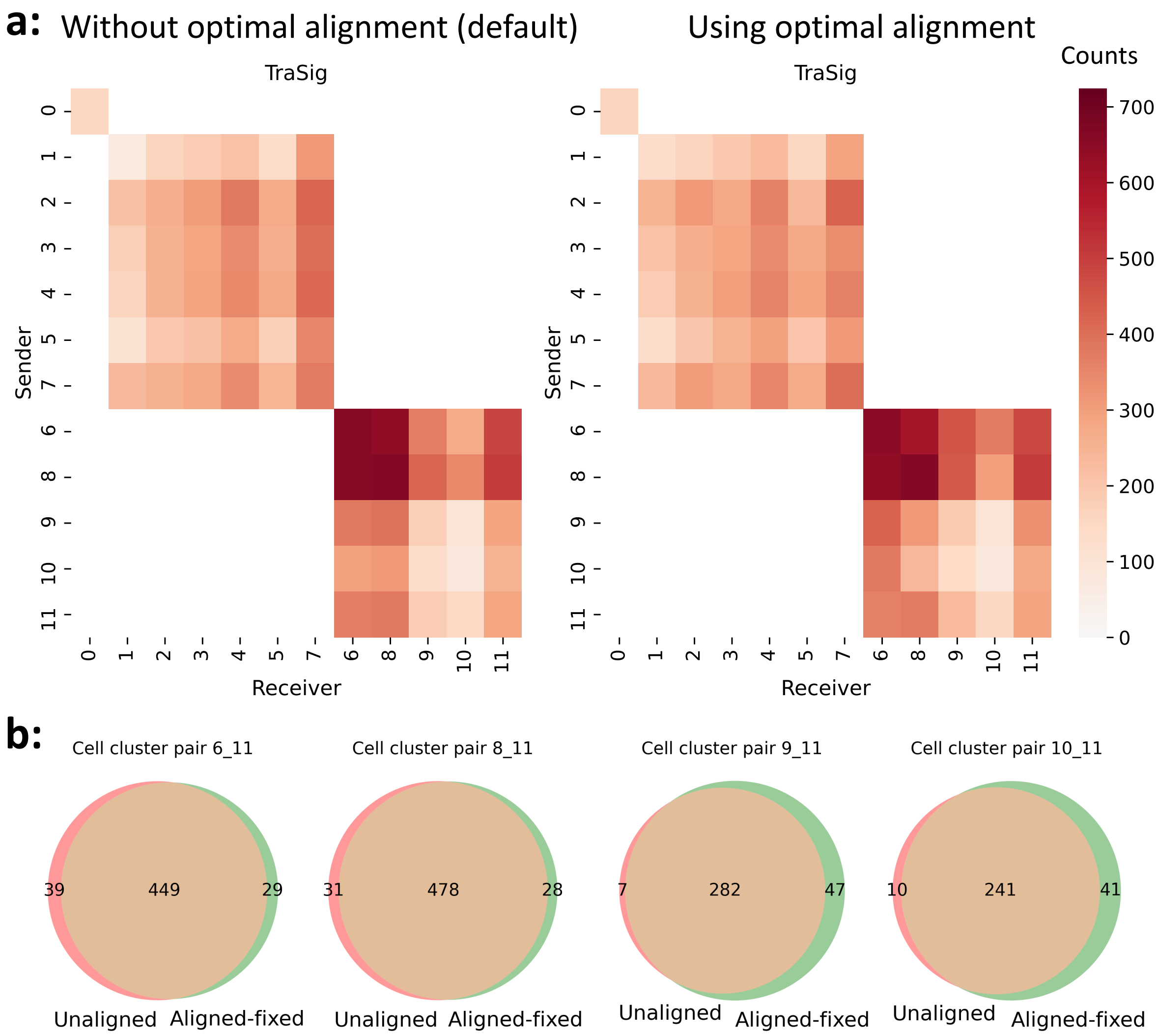


Figure S12: We compare TraSig's inference results on the liver organoid data between the default option without using optimal alignment and when optimal alignment is applied. (a) Number of significant ligand-receptor pairs identified between each edges (clusters) pair. The overall interaction patterns when using or not using optimal alignment are similar. Both heatmaps use the same anchor values (minimum: 0, maximum: 724) for the colormap. (b) Venn diagrams for the overlap in the identified ligand-receptor pairs for some representative edges (clusters) pairs. Unaligned: default; Aligned-fixed: using optimal alignment. Note here we performed 10,000 permutations instead of 100,000 permutations as used for the other results.

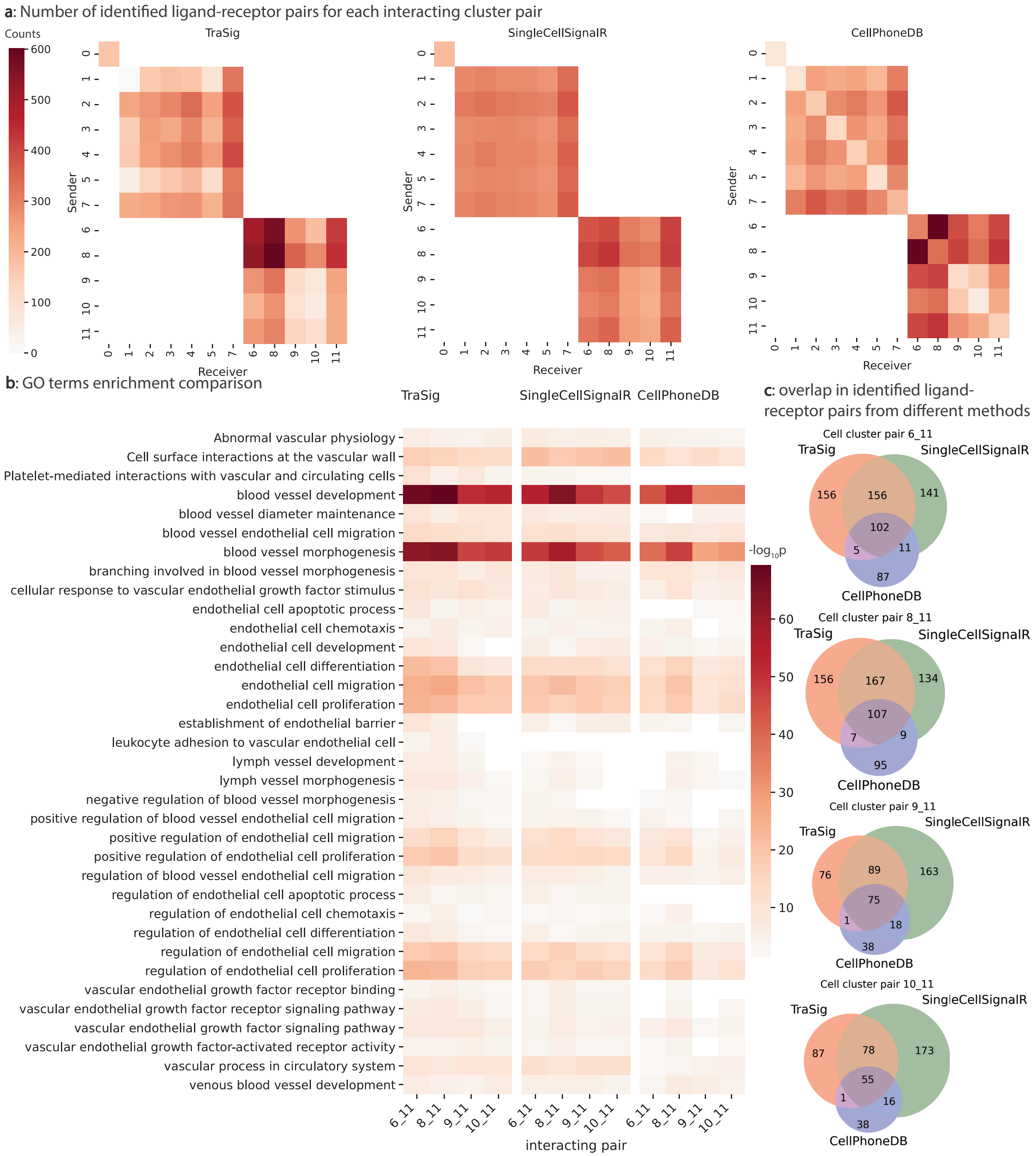


Figure S13: Results from comparing TraSig with SingleCellSignalR and CellPhoneDB using another database [7] on the liver organoid data. (a) Heatmaps for scores assigned by the three different methods for all cluster pairs representing cells sampled at the same time. (b) $-\log_{10}$ p-value for enriched GO terms related to endothelial cells and vascular development. (c) Venn diagrams for the overlap in identified ligand-receptor pairs among the three methods. The overlap between TraSig and SingleCellSignalR is high though roughly 50% of the identified pairs by each method are not identified by the other.
